# Arbuscular mycorrhizal fungi influence host infection during epidemics in a wild plant pathosystem

**DOI:** 10.1101/2021.09.28.462160

**Authors:** Jenalle L. Eck, Minna-Maarit Kytöviita, Anna-Liisa Laine

## Abstract

While pathogenic and mutualistic microbes are ubiquitous across ecosystems and often co-occur within hosts, how they interact to determine patterns of disease in genetically diverse wild populations is unknown.
To test whether microbial mutualists provide protection against pathogens, and whether this varies among host genotypes, we conducted a field experiment in three naturally-occurring epidemics of a fungal pathogen, *Podosphaera plantaginis*, infecting a host plant, *Plantago lanceolata*, in the Åland Islands, Finland. In each population, we collected epidemiological data on experimental plants from six allopatric populations that had been inoculated with a mixture of mutualistic arbuscular mycorrhizal fungi or a non-mycorrhizal control.
Inoculation with arbuscular mycorrhizal fungi increased growth in plants from every population, but also increased host infection rate. Mycorrhizal effects on disease severity varied among host genotypes and strengthened over time during the epidemic. Host genotypes that were more susceptible to the pathogen received stronger protective effects from inoculation.
Our results show that arbuscular mycorrhizal fungi introduce both benefits and risks to host plants, and shift patterns of infection in host populations under pathogen attack. Understanding how mutualists alter host susceptibility to disease will be important for predicting infection outcomes in ecological communities and in agriculture.

**Plain Language Summary:** Beneficial, ‘mycorrhizal’ fungi in roots help plants grow and may protect them from diseases caused by pathogenic microbes. This study shows that arbuscular mycorrhizal fungi can influence patterns of plant disease during pathogen outbreaks in a natural landscape.

## Introduction

Protective symbionts – species that provide defensive benefits to their hosts – help determine the outcome of species interactions and, thus, shape ecological and evolutionary dynamics between hosts and parasites (Brownlie & Johnson, 2009; May & Nelson, 2014; King *et al*., 2016; Sochard *et al*., 2020). Despite their importance, ecological studies examining the role of protective symbionts in influencing host-parasite interactions in natural populations and communities are rare (Oliver *et al*., 2014; Hafer-Hahmann & Vorburger, 2021). Protection against infectious disease by mutualistic microbes, such as mycorrhizal fungi, has been demonstrated under controlled laboratory conditions in several economically important agricultural plant species (Norman *et al*., 1996; Pozo *et al*., 2002; Li *et al*., 2010; Song *et al*., 2015; Berdeni *et al*., 2018). However, it has not been verified whether protective symbionts mediate infection under natural epidemics, which are characterized by repeated pathogen encounters, as well as by environmental and genotypic diversity in both hosts and pathogens (Newsham *et al*., 1995). Mycorrhizal associations are widespread among terrestrial plants (Öpik *et al*., 2006; van der Heijden *et al*., 2008), and have important impacts on plant fitness and population dynamics (Barea *et al*., 2002; Koide & Dickie, 2002), community composition (Hartnett & Wilson, 2002) and ecosystem functioning (Rillig, 2004). Understanding how mycorrhizal fungi and other mutualists influence patterns of plant disease is essential given that disease is a major factor shaping the abundance, diversity, and distribution of species in plant communities (Bever *et al*., 2015) and affecting food production (Johansson *et al*., 2004; Gosling *et al*., 2006; Pretty *et al*., 2011; Hohmann & Messmer, 2017). While both plant-associated pathogenic and mutualistic microbes are ubiquitous across ecosystems, how they interact to determine disease risk in natural, genetically diverse populations is not known.

Mycorrhizal fungi produce a suite of growth, nutritional, and/or defensive effects that may help protect plants from co-occurring antagonists, such as pathogenic microbes and herbivores (Delavaux *et al*., 2017). Association with mycorrhizal fungi often improves plant nutrient and water uptake (Smith & Read, 2008), though there is both intra- and interspecific variation among plants in their ability to form and benefit from mycorrhizal associations (Thrall *et al*., 2011; Rasmussen *et al*., 2019). Increases in host size and nutritional status due to mycorrhizal association can improve host tolerance to parasites and abiotic stress (Azcón-Aguilar & Barea, 1996). Arbuscular mycorrhizal fungi can also influence host defenses directly, by upregulating defense gene expression in their host plants (Azcón-Aguilar & Barea, 1996; Pozo & Azcón-Aguilar, 2007; Jung *et al*., 2012). This form of protection, known as defense priming, allows a more efficient activation of defense mechanisms in response to attack by potential enemies, and has been shown to reduce the negative effects of interactions with a wide range of antagonist species (Jung *et al*., 2012; Delavaux *et al*., 2017). Though mycorrhizal associations occur belowground (at the root-soil interface), the induced resistance response in the host is systemic (Cameron *et al.*, 2013), meaning that even strictly aboveground parasites may be affected (Koricheva *et al*., 2009). It is unclear how often mycorrhizal growth and defensive benefits are conferred together in hosts, and how they operate simultaneously to determine the incidence and outcome of interactions between hosts and parasites.

Empirical studies in controlled environments have shown that host association with mutualists may also present risks that may influence infection dynamics (Polin *et al*., 2014). For example, ecological costs may occur when combinations of host and mutualist species or genotypes are mismatched (Klironomos, 2003; Hoeksema *et al*., 2010), resulting in inefficient mutualisms that fail to convert host resources into growth or defensive benefits (Johnson *et al*., 1997; Jones & Smith, 2004; Grman, 2012). Furthermore, unfavorable abiotic conditions may reduce or negate potential mutualist benefits (Hoeksema *et al*., 2010; Qu *et al*., 2021). In addition, defensive benefits from mutualists may not be effective against all parasite species (e.g., depending on their life history) (Pozo & Azcón-Aguilar, 2007), or durable to changes in parasite traits and/or composition in the environment. Finally, mutualists could also affect patterns of host infection indirectly, e.g., via changes in host size that influence parasite contact rates. Hence, the potential ecological risks and/or benefits of mutualist association in the presence of parasites may depend on host genotype and vary among or within host populations and environments; however, this has remained largely unexplored in natural populations.

In addition, it is unclear how mutualism-derived protection acts alongside innate host resistance to determine the outcome of host-pathogen interactions. Host genetic resistance can vary widely among and within natural populations (Salvaudon *et al*., 2008, Laine *et al*., 2011) - potentially due to costs associated with its maintenance (Brown, 2003; Susi & Laine, 2015) - and may be under a different set of selection pressures than mutualism (Thompson, 1994). Whether symbiosis presents benefits or costs to host individuals and populations could depend on their level of resistance. In resistant hosts, resources provided to mutualists in return for defensive benefits could represent an unnecessary metabolic cost. However, in susceptible hosts, mutualist-derived protection could compensate for lack of genetic resistance, presenting a viable alternate strategy for coping with pathogens. Mutualism-derived protection could be especially beneficial when genetic resistance is costly to maintain, or when disease is ephemeral, as it often is in natural populations (Burdon & Thrall, 2014). How much of host resistance is derived from innate genetic defenses versus derived from protective symbionts (e.g., defense priming, or improved pathogen tolerance), and whether these types of defenses are linked in different species combinations and environmental contexts, remains to be seen.

Despite general recognition for the impact of mycorrhizal fungi on plant fitness, how mycorrhizal association may impact host infection – and to what extent this varies among plant genotypes and populations – is poorly understood in natural populations. To examine this, we conducted a field experiment to test whether inoculation with arbuscular mycorrhizal fungi affects infection dynamics by a fungal pathogen in a shared host under natural epidemic conditions. Specifically, we ask: 1) Do growth effects due to inoculation with arbuscular mycorrhizal fungi vary among host populations and maternal genotypes? 2) Does inoculation with arbuscular mycorrhizal fungi influence host infection rate? 3) Upon infection, does prior inoculation with arbuscular mycorrhizal fungi affect disease severity? 4) Are growth and defensive effects from inoculation with arbuscular mycorrhizal fungi linked in host genotypes? and 5) Are disease susceptibility and defensive effects due to inoculation with arbuscular mycorrhizal fungi linked in host genotypes? To answer these questions within an ecologically-relevant context, we placed mycorrhizal-inoculated and non-mycorrhizal-inoculated plants in wild host populations during a natural pathogen epidemic. The experiment was conducted in the long-term Åland Islands study site (Finland), where infection by a fungal pathogen (powdery mildew) has been surveyed on a large host population network of *Plantago lanceolata* L. (Plantaginaceae) since 2001 (Jousimo *et al*., 2014). From prior studies in this pathosystem, we know that several factors, such as spatial context (Laine, 2006; Soubeyrand *et al*., 2009; Jousimo *et al*., 2014), pathogen genetic diversity (Eck *et al*., 2022), local adaptation of pathogen strains to sympatric host populations (Laine, 2005; Laine, 2007a), and abiotic conditions (Laine, 2007b; Laine, 2008) are all critical in determining infection, but the impact of mutualistic interactions on infection dynamics has not been determined. To our knowledge, this is the first study of the impact of mycorrhizal fungi on infection by a plant pathogen under natural epidemics and across different host populations and genotypes.

## Materials and Methods

### Study system

Our study is focused on a fungal pathogen, *Podosphaera plantaginis* (Castagne) U. Braun & S. Takam., infecting *Plantago lanceolata* L. (Plantaginaceae) in the Åland Islands (60°08’53” N, 19°47’18” E), Finland. We carried out an experiment in three natural *Pl. lanceolata* populations that are part of a network of > 4000 mapped populations (Hanski, 1999). These populations have been surveyed for infection by *Po. plantaginis* since 2001 (Laine & Hanski, 2006). *Plantago lanceolata* (ribwort plantain) is native to Åland and much of Eurasia; it occurs mainly in small meadows and disturbed areas in Åland. It is monoecious, self-incompatible, and reproduces either sexually (via wind-dispersed pollen and seeds; Bos, 1992) or asexually (via clonally-produced side-rosettes) (Sagar & Harper, 1964). *Plantago lanceolata* associates commonly with mycorrhizal fungi and has been found in association with a wide variety of mycorrhizal species in Åland (Rasmussen *et al*., 2018; Eck *et al*., unpublished data).

*Podosphaera plantaginis*, a powdery mildew fungus (order Erysiphales), is an obligate biotroph of foliar tissue, and is host-specific in Åland, infecting only *Pl. lanceolata* (Laine, 2004). Fungal hyphae grow on the surface of *Pl. lanceolata* leaves, producing localized infections that mitigate host growth and reproduction (Bushnell, 2002), and may lead to mortality in the presence of abiotic stress, such as drought (Laine, 2004; Susi & Laine, 2015). Infections are transmitted via asexually-produced, wind-dispersed spores (conidia) that are produced cyclically (c. every two wks) throughout an epidemic season (c. June to September in Åland) (Ovaskainen & Laine, 2006). Resting structures (chasmothecia), produced via haploid selfing, or outcrossing between strains (Tollenaere & Laine, 2013), allow the pathogen to overwinter (Tack & Laine, 2014). In Åland, *Po. plantaginis* persists as a metapopulation, with frequent colonization and extinction events (Jousimo *et al*., 2014). Resistance in *Pl. lanceolata* against *Po. plantaginis* is strain-specific (Laine, 2004; Laine, 2007a). Previous studies have demonstrated high levels of variation in resistance within and among host populations (Laine, 2004; Laine 2007a, Jousimo *et al*. 2014).

### Mycorrhizal inoculation of experimental plants

To measure the effects of association with arbuscular mycorrhizal fungi in genetically diverse host plants, seeds were collected from six geographically variable populations of *Pl. lanceolata* in the Åland Islands (Fig. 1A) in August 2007. Seeds were collected from five haphazardly chosen maternal plants in each population and were stored separately in paper envelopes (seeds from one maternal plant are half or full siblings). In April 2008, seeds from each of the 30 maternal plants were planted in separate 6 × 6 × 7 cm pots in a glasshouse at the University of Oulu (Oulu, Finland) in sterilized sand. Ten healthy two-week-old seedlings from each maternal genotype were transferred to individual pots (one seeding per 6 × 6 × 7 cm pot) filled with experimental substrate (a 5: 4: 1 mixture of heat-sterilized sand: heat-sterilized garden soil: perlite, combined with 1 g of bone meal and 3 g of dolomite per liter substrate). At the time of transfer, five of the seedlings from each maternal genotype were inoculated with mycorrhizal fungi, while the other five received a non-mycorrhizal control treatment. The mycorrhizal inoculum consisted of a mixture of spores of three arbuscular mycorrhizal fungal species native to Finland (*Glomus hoi*, *Claroideoglomus clareoideum*, and *Gl. mosseae (BEG 29)*), as *Pl. lanceolata* is colonized naturally by several mycorrhizal symbionts (Johnson *et al*., 2003). Allopatric isolates of each species (originating from central Finland, 61º10 N, 24º40 E, rather than Åland) were used so as not to confound the experiment with potential local adaptation between mycorrhizal fungi and their host populations (Hoeksema *et al*., 2010). Spore inocula were produced by growing the mycorrhizal species with non-experimental *Pl. lanceolata* in a soil substrate identical to the experimental one. Spores were washed out of the non-experimental substrate with water, and 15 spores of each species (45 spores in total) were pipetted on to the roots of each seedling in the mycorrhizal treatment in 2 ml of water (6 ml in total); seedlings in the non-mycorrhizal control treatment received 2 ml of filtered spore washing water from each fungal species (6 ml in total). Hereafter, plants in the mycorrhizal-inoculated treatment are abbreviated as ‘AMF’ (i.e., arbuscular mycorrhizal fungi), and plants in the non-mycorrhizal control treatment as ‘NM’ (i.e., non-mycorrhizal). All experimental seedlings were also inoculated with bacteria at this time: bacteria were filtered from a soil mix collected from the six Åland seed source populations to restore the native, non-mycorrhizal soil microbial community. Seedlings were fertilized weekly with a dilute nitrogen-based solution. Natural light in the glasshouse was supplemented with Osram HQI lamps to provide a photoperiod of 18 h: 6 h light : dark. At 6 wks of age (4 wks post-inoculation), the plants were moved to an outdoor area at the University of Oulu and grown on tables under a transparent plastic roof for an additional six weeks, to acclimatize to field conditions (as *Pl. lanceolata* and *Po. plantaginis* do not inhabit this region, infection at this stage was highly unlikely).

**Figure 1:**
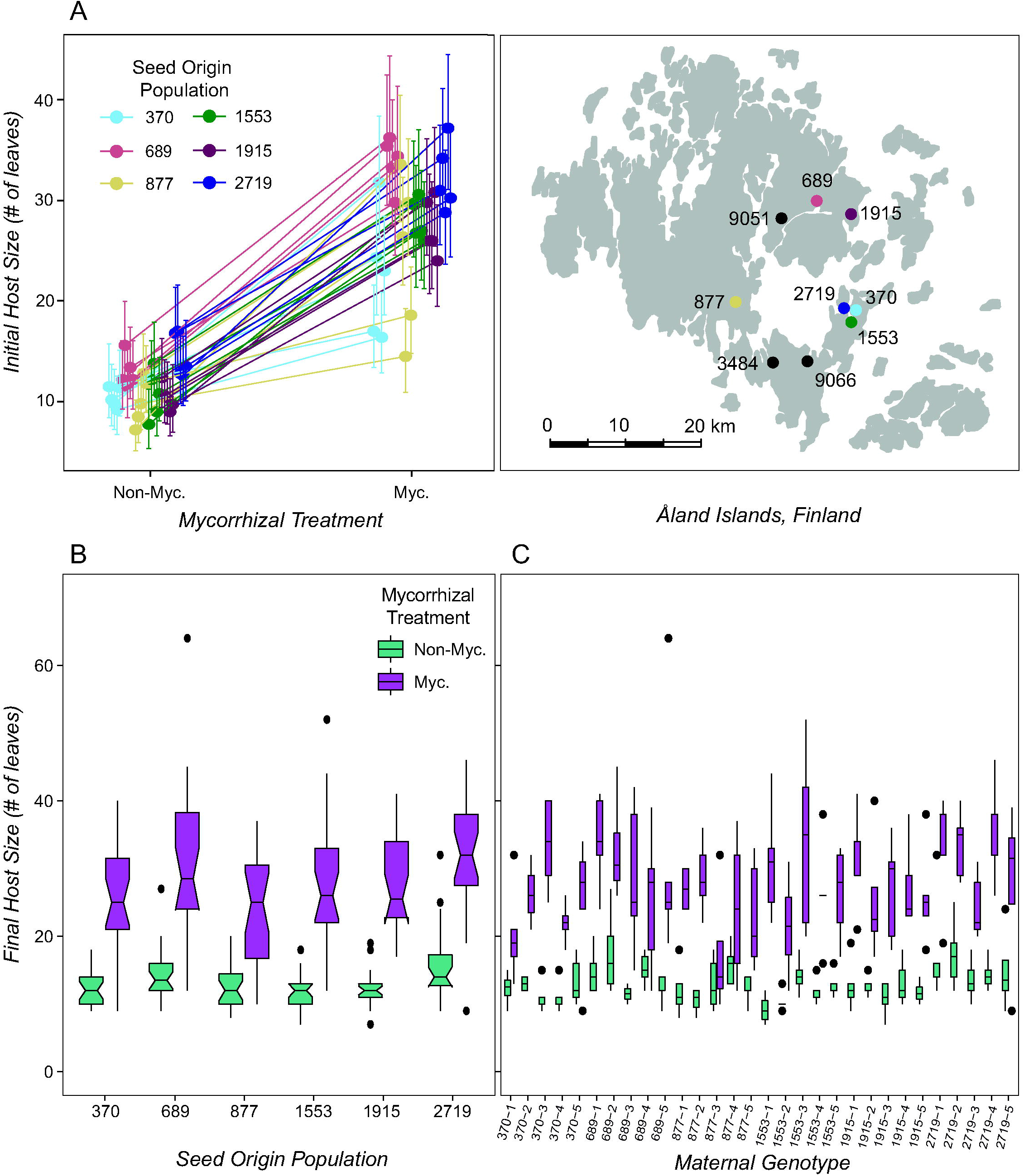
Inoculation with arbuscular mycorrhizal fungi produced growth benefits in experimental plants from most genetic origins. Before exposure to *Podosphaera plantaginis* in field conditions, the magnitude of the growth benefits in experimental *Plantago lanceolata* plants following inoculation with a mixture of three mycorrhizal fungal species varied among thirty host maternal genotypes (*Panel A*; Supporting Information Table S2; MYC × GEN: *p* < 0.001, *n* = 287 plants). In *A*, colored dots represent seed origin populations and black dots represent field epidemic populations. After pathogen exposure in the field epidemic experiment, host growth continued to be linked to mycorrhizal inoculation (*Panels B & C*; Supporting Information Tables S3 & S4; MYC: *p* < 0.001), seed origin population (*Panel B*; Table S3; POP: *p* < 0.001), and maternal genotype (*Panel D*; Supporting Information Table S4; GEN: *p* < 0.001), but growth benefits no longer varied among maternal genotypes. In *A*, error bars represent a 95% confidence interval. In *B* & *C*, box notches represent a 95 % confidence interval for comparing medians, box hinges correspond to the 1^st^ and 3^rd^ quartiles, while box whiskers extend to the largest and smallest value no further than 1.5 × the interquartile range from the hinges.

### Natural epidemic field experiment

Ten weeks post-inoculation (mid-July 2008), 288 healthy 12-wk-old experimental plants were placed in to three naturally-occurring populations of *Pl. lanceolata*, infected by populations of *Po. plantaginis*, in Åland to gain infections naturally from the surrounding epidemic. The three field epidemic populations were allopatric to the six seed origin populations (Fig. 1A). Thus, they represent common garden field sites, in which the host genotypes are not locally adapted to the local environmental conditions or to the local pathogen population. Presence of *Po. plantaginis* at the sites was confirmed by surveying the populations for visible signs of infection before placement of the experimental plants. Each field site received 96 experimental plants: 16 plants from each of the six seed origin populations (eight AMF plants and eight NM plants). Within these subgroups, the five maternal genotypes within each seed origin population were allocated as evenly as possible to the three field sites; the individuals placed in each site were selected at random (full, pairwise replication of every maternal genotype × mycorrhizal treatment combination within every field site was not possible due to lack of plants). At each field site, experimental plants were randomly fitted in to one of 96 plastic containers (14 × 10.5 × 4.5 cm) that were placed on the ground in a random array in the proximity of naturally-infected, wild *Pl. lanceolata* individuals inhabiting the field population. The containers prevented contact between the bottom of the pots and the soil, reducing the likelihood that the experimental plants would acquire soil microbes from the environment.

At the time of placement in the field epidemic populations, we measured the initial size of each experimental plant: the total number of leaves, as well as the length and width (in cm) of the longest leaf were recorded. Plants were considered infected if any leaf showed powdery, white spot(s) (i.e., characteristic signs of infection with *Po. plantaginis* in this region; Supporting Information Fig. S1) forming a blotch or lesion of any size upon visual inspection (all leaves were inspected for signs of infection). All experimental plants were uninfected (i.e., had zero leaves infected by *Po. plantaginis*) at the beginning of the experiment. Infection surveys were then conducted every three days at each field site, on a rotating survey schedule (1^st^ day field population 9051, 2^nd^ day field population 3484, and 3^rd^ day field population 9066). During each infection survey, the number of healthy leaves and leaves infected by *Po. plantaginis* on each plant were counted (leaves that withered during the experiment were not counted). The plants were also re-randomized into a new position in the experimental array (this was done to minimize the effect of spatial positioning, with respect to distance to infected individuals and prevailing wind direction, on infection rate and severity), and were watered if necessary. Seven infection surveys were conducted in each field population. At the end of the experiment (in late August 2008) a haphazardly-selected subset of 21 AMF plants and 20 NM plants were harvested, and their oven-dried aboveground biomass was measured. We created a metric approximating the leaf area of each plant at the beginning of the experiment by first applying the length and width of the plant’s longest leaf to the equation yielding the area of an oval, then multiplying the area of the longest leaf by the total number of leaves on the plant. Heavy rains during the 4^th^ and 5^th^ surveys resulted in missing infection data for some field populations (identifying symptoms caused by *Po. plantaginis* on wet leaves is challenging) and may have influenced infection in the 6^th^ and 7^th^ surveys in all populations (as heavy rain washes spores away from infected leaf tissue and damages spore viability; Sivapalan, 1993). Because of this, we focus here on infection data from the 3^rd^ survey (coinciding with peak infection rates and the completion of one 14-day initial pathogen infection and reproduction cycle), as well as the final, 7^th^ survey at the end of the experiment (coinciding with re-infections because of pathogen reproduction and incorporating variable abiotic conditions). Using data from these two surveys (peak epidemic and end-of-experiment), we quantified the infection status (0 / 1, infected or uninfected) and calculated the infection severity (i.e., the proportion of infected leaves) of each experimental plant. Plants were considered infected if one or more leaves showed signs of infection. One plant that died during the experiment was excluded from all analyses.

### Statistical methods

#### Do growth effects due to inoculation with arbuscular mycorrhizal fungi vary among host populations and maternal genotypes?

To test whether host growth was explained by inoculation with arbuscular mycorrhizal fungi, host genetic origin, or an interaction between these factors, we built a series of generalized linear models (GLMs). All statistical tests were conducted in the R statistical environment (R Core Team, 2021). Negative binomial GLMs were constructed with the MASS package counter overdispersion (Venables & Ripley, 2002). To explore the effect of genetic origin on host growth at two levels of biological organization, seed origin population (abbreviation: POP) and maternal genotype (abbreviation: GEN) were included as explanatory factors in separate models. Host growth was modeled before epidemic exposure (at the time of placement in the field experiment) and post-epidemic exposure (during the last survey of the experiment) using the number of leaves on each host. When modeling initial host size, the explanatory factors were mycorrhizal treatment (abbreviation: MYC), POP/GEN, and their interaction. When modeling final host size, field epidemic population (abbreviation: SITE), initial host size (abbreviation: SIZE), and a four-way interaction term between all factors were also included as explanatory factors whenever possible (without compromising model fit). We also tested whether a subset of AMF and NM plants varied in aboveground dry biomass at the end of the experiment using a Mann-Whitney U test to counter non-normality in the data. Throughout our study, non-significant interaction terms were removed from all final models. To address non-independence of the mycorrhizal treatment and host size variables during the field experiment, all models including size were also tested with this variable omitted, and the results were compared.

#### Does inoculation with arbuscular mycorrhizal fungi influence host infection rate?

To test whether host infection rate upon exposure to field epidemics is explained by inoculation with arbuscular mycorrhizal fungi, host genetic origin, field epidemic population, or an interaction between these factors, we built a series of generalized logistic regression models. Host infection status was used as the binomial response variable in each model (with family set to quasibinomial to counter overdispersion). Infection data were analyzed at the peak of the epidemic and at the end of the experiment. In each model, MYC, POP/GEN, SITE (and SIZE at the corresponding time point, in applicable models) were included as explanatory factors, as well as all interaction terms.

#### Upon infection, does prior inoculation with arbuscular mycorrhizal fungi affect disease severity?

To test whether disease severity in infected hosts is influenced by prior inoculation with arbuscular mycorrhizal fungi, host genetic origin, field epidemic population, or an interaction between these factors, we built a series of generalized logistic regression models. These models explored two measures of disease severity (i.e., the proportion of leaves infected and the number of infected leaves in infected individuals) at two time points (i.e., at the peak of the epidemic and at the end of the experiment). In each model, weights were set to the total number of leaves on the individual; family was set to binomial in models of the proportion of leaves infected, and to quasipoisson in models of the number of leaves infected (to counter overdispersion). In each model, MYC, POP/GEN, SITE (and SIZE at the corresponding time point, in applicable models) were included as explanatory factors, as well as all interaction terms.

#### Are growth and defensive effects from mycorrhizal inoculation linked in host genotypes?

To test for a relationship between growth and defensive effects following inoculation with arbuscular mycorrhizal fungi in the maternal genotypes, we first used the negative binomial GLM examining differences in host size (i.e., the model built in the first section of the statistical methods) to calculate the estimated effect of mycorrhizal inoculation on host growth in each maternal genotype. Host growth effects were estimated at the beginning of the experiment to avoid the effects of pathogen exposure and variable field conditions. Effect sizes were obtained using the lsmeans package (Lenth, 2016) and were averaged over the levels of field epidemic population. Second, we used the generalized logistic regression model examining differences in the number of infected leaves in infected hosts (i.e., the model built in the third section of the statistical methods) to calculate the estimated effect of mycorrhizal inoculation on host defense in each maternal genotype. Defensive effects were quantified at the epidemic peak to capture the highest infection levels. We then built a linear regression model linking defense effects (as the response variable) and growth effects in each genotype.

#### Are disease susceptibility and defensive effects from mycorrhizal inoculation linked in host genotypes?

We tested for a relationship between host disease susceptibility and defensive effects following inoculation with arbuscular mycorrhizal fungi in the host genotypes. First, we quantified disease susceptibility in each genotype in the absence of the mutualist (i.e., in NM plants only) using the estimated coefficients from the model examining the number of infected leaves at the epidemic peak (i.e., the same model as used in the section above). Genotype-level coefficients were obtained using the lsmeans package (Lenth, 2016) and were averaged over the levels of field epidemic population. Second, defensive effects due to inoculation with mycorrhizal fungi were quantified for each maternal genotype as the effect of mycorrhizal inoculation from these same models (i.e., as the change in the coefficient when the genotype was inoculated). Finally, we built a linear regression model linking defense effect sizes (as the response variable) and disease susceptibility in each genotype.

## Results

### Do growth effects due to mycorrhizal inoculation vary among host populations and maternal genotypes?

Inoculation with arbuscular mycorrhizal fungi produced growth benefits in experimental plants from nearly every host genetic origin. Before pathogen exposure in field conditions, host growth was determined by mycorrhizal inoculation and seed origin population (Supporting Information Fig. S2 & Table S1; MYC: *p* < 0.001, POP: *p* < 0.001, *n* = 287 plants), as well as by an interaction between mycorrhizal inoculation and host maternal genotype (Fig. 1A & Supporting Information Table S1; MYC × GEN: *p* = 0.002). At the end of field epidemic experiment, mycorrhizal inoculation (Fig. 1B – 1C & Supporting Information Tables S2 – S3; MYC: *p* < 0.001), seed origin population (Fig. 1B & Supporting Information Table S2; POP: *p* < 0.001), and maternal genotype (Fig. 1C & Supporting Information Table S3; GEN: *p* < 0.001) continued to be tightly linked to host growth, but the interaction between mycorrhizal inoculation and maternal genotype had disappeared. Host initial size also predicted of host final size (Supporting Information Table S2; SIZE: *p* < 0.001). Mycorrhizal inoculation also increased the aboveground dry biomass of plants at the end of the experiment (Supporting Information Fig. S3 & Table S4; MYC: *p* < 0.001, *W* = 11, *n* = 41 plants).

### Does inoculation with mycorrhizal fungi influence host infection rate?

Upon pathogen exposure in the field epidemics, host infection rate was influenced by several factors whose importance shifted over time. Host infection rates reached 73 % at the peak of the epidemic but fell to 34 % by the end of the experiment (Fig. 2A). Host infection rate also varied among the three field epidemic populations, with the effect of field site strengthening over time (Figs. 2B – 2D & Supporting Information Tables S5 – S8; SITE (peak of epidemic): *p* = 0.03 (POP) – 0.01 (GEN); end of experiment: *p* < 0.001 (POP/GEN). Though inoculation with arbuscular mycorrhizal fungi marginally influenced host infection rate during the peak of the epidemic (Fig. 2C & Supporting Information Tables S5 – S6; MYC: *p* = 0.09 (POP) - 0.11 (GEN), *n* = 287 plants), at the end of experiment AMF plants were slightly more likely to be infected than NM plants (Fig. 2D & Supporting Information Tables S7 – S8; MYC: *p* = 0.04 (POP) – 0.05 (GEN), *n* = 286 plants). The effect of mycorrhizal inoculation on host infection rates was weaker than the effect on host growth. Host size also influenced host infection at the end of the experiment: larger plants were more likely to be infected (Supporting Information Tables S7 – S8; SIZE: *p* = 0.06 (POP) – 0.04 (GEN)).

**Figure 2:**
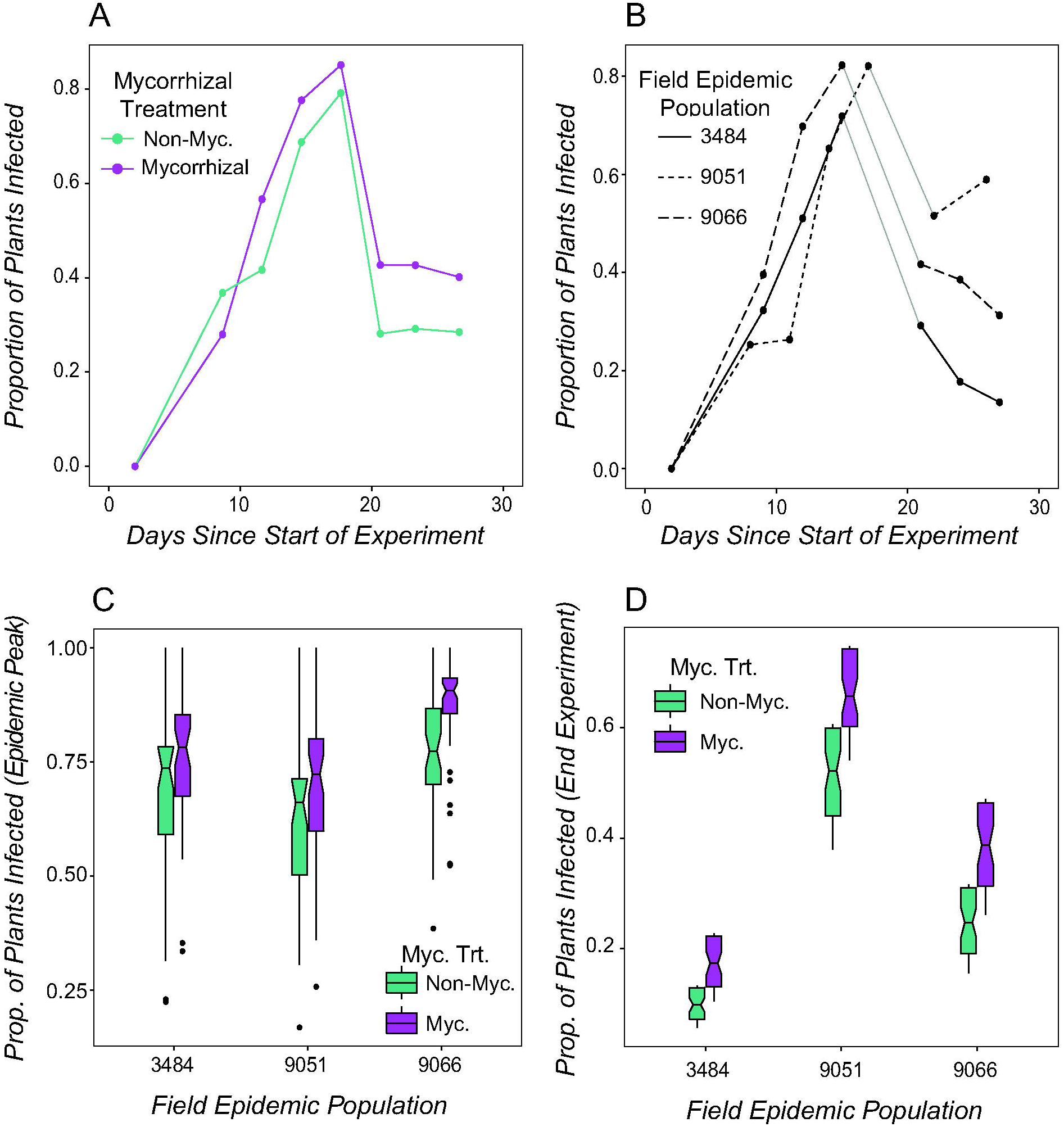
Host infection rate was increased in mycorrhizal-inoculated plants and varied among field epidemic populations. Infection rates by *Podosphaera plantaginis* in experimental *Plantago lanceolata* plants increased over time in each mycorrhizal treatment (*Panel A*) and field epidemic population (*Panel B*) until the peak of the epidemic, but fell by the end of the experiment. Though host infection rate was similar between mycorrhizal treatments at the peak of the epidemic (*Panel C*; Supporting Information Tables S5 - S6), at the end of the experiment host infection rates were increased in mycorrhizal-inoculated plants relative to NM plants in all three field epidemic populations (Supporting Information Tables S7 – S8; MYC: *p* = 0.04, *n* = 286 plants). Host infection rates also varied among the three field epidemic populations throughout the experiment (*Panels C* & *D*; Supporting Information Tables S5 – S8; SITE: *p* = 0.01 (peak), *p* < 0.001 (end), *n* = 286 plants). In *B*, missing data due to heavy rains on days c. 15 - 20 are indicated by light grey solid lines. In *C* & *D*, predicted values resulting from generalized logistic models are plotted; box notches represent a 95 % confidence interval for comparing group medians, box hinges correspond to the 1^st^ and 3^rd^ quartiles, while box whiskers extend to the largest and smallest value no further than 1.5 × the interquartile range from the hinges.

### Upon infection, does prior inoculation with mycorrhizal fungi affect disease severity?

Among infected individuals, disease severity was influenced by several factors, including mycorrhizal inoculation and host genetic origin. AMF plants had lower proportions of their leaves infected relative to NM plants throughout the experiment (Figs. 3A & 3B, Supporting Information Tables S9 – S12; MYC: *p* < 0.001, *n* = 210 plants (peak epidemic); *p* = 0.003, *n* = 98 plants (end experiment)). The proportion of leaves infected also varied among seed origin populations (Fig. 3A & Supporting Information Table S9; POP: *p* = 0.004) and maternal genotypes (Supporting Information Table S10; GEN: *p* = 0.02) during the peak of the epidemic but was similar among host genetic origins by the end of the experiment (Supporting Information Tables S11 – S12). At the epidemic peak, mycorrhizal inoculation and maternal genotype interacted to determine the number of infected leaves on infected hosts (Fig. 3C & Supporting Information Table S14; MYC × GEN: *p* = 0.05), with both positive and negative effects occurring. By the end of the experiment, AMF plants had higher numbers of infected leaves than NM plants (Fig. 3D & Supporting Information Table S16; MYC: *p* = 0.04). Like the effect on host infection rates, the effect of mycorrhizal inoculation on disease severity was weaker than the effect on host growth. Field epidemic population also influenced the number of infected leaves in hosts at the end of the experiment (Fig. 3D & Supporting Information Table S15; SITE: *p* = 0.02). Host size predicted the proportion of infected leaves at the peak of the epidemic (Supporting Information Tables S9 – S10; SIZE: *p* < 0.001) and the number of infected leaves throughout the experiment (Supporting Information Tables S13 – S16; SIZE: *p* < 0.01).

**Figure 3:**
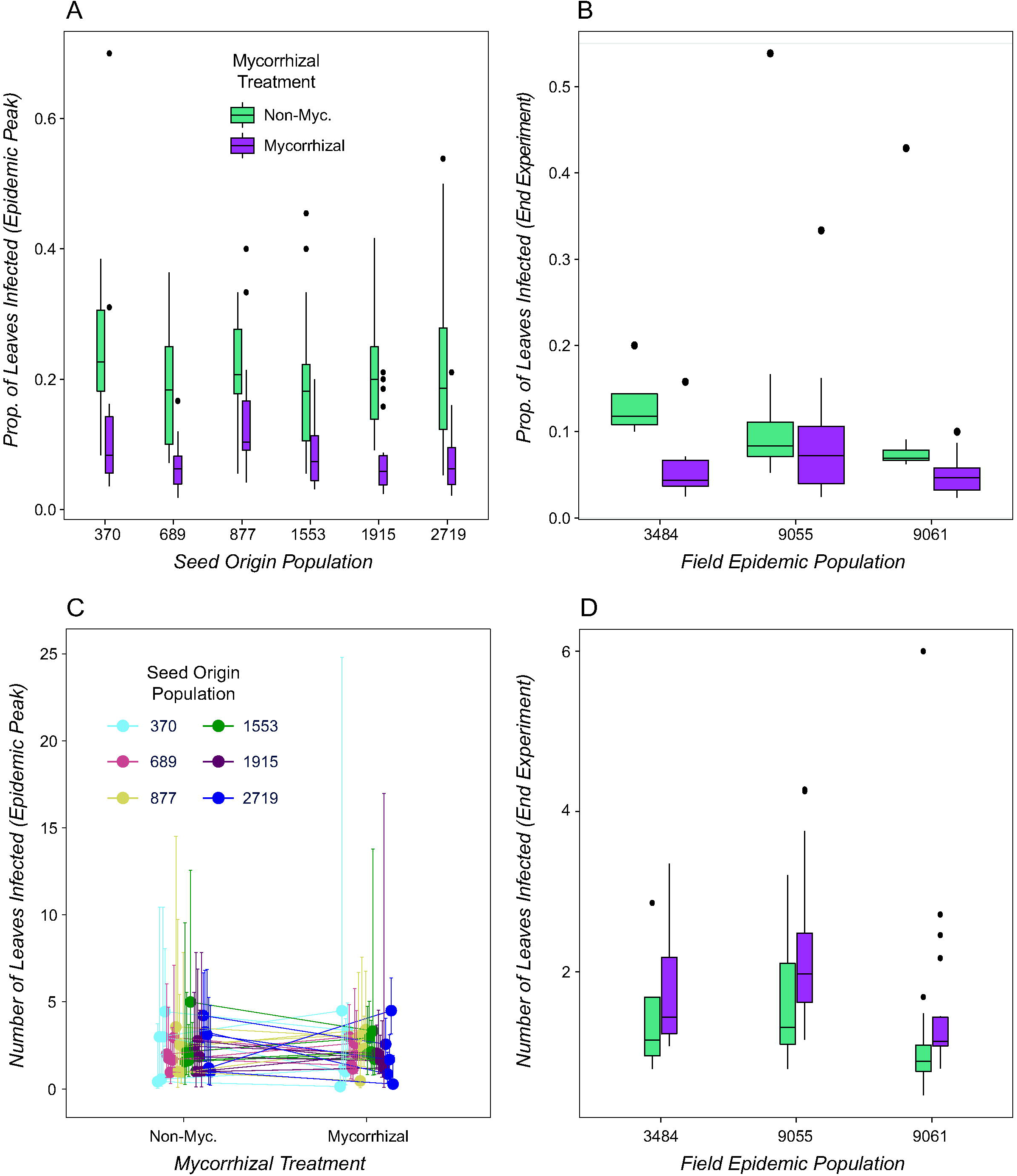
Among infected plants, disease severity varied among mycorrhizal treatments, seed origin populations, and field epidemic sites. Prior inoculation with a three-species mixture of arbuscular mycorrhizal fungi was linked to reductions in the proportion of leaves infected by *Podosphaera plantaginis* in infected individuals of *Plantago lanceolata*, both at the peak of the epidemic (*Panel A*; Supporting Information Tables S9 – S10; MYC: *p* < 0.001, *n* = 210 plants) and at the end of the experiment (*Panel B*; Supporting Information Tables S11 – S12; MYC: *p* = 0.003, *n* = 98 plants). Mycorrhizal inoculation and maternal genotype also interacted to determine the number of infected leaves on infected plants at the peak of the epidemic, with both negative and positive effects of mycorrhizae on disease (*Panel C*; Supporting Information Table S14, MYC × GEN: *p* = 0.05). At the end of the experiment, infected AMF plants had more infected leaves than infected NM plants (*Panel D*; Supporting Information Table S16; MYC: *p* = 0.04). Seed origin population (*Panel A*; Supporting Information Table S9; POP: *p* = 0.004) and field epidemic population (*Panel D*; Supporting Information Tables S14 – S16; SITE: *p* < 0.02) also influenced host disease severity. In *A – B & D*, predicted values resulting from generalized logistic models are plotted; box hinges correspond to the 1^st^ and 3^rd^ quartiles, while box whiskers extend to the largest and smallest value no further than 1.5 × the interquartile range from the hinges. In *C*, error bars represent a 95 % confidence interval.

### Are growth and defensive effects from mycorrhizal inoculation linked in host genotypes?

There was a marginally significant relationship between the growth and defensive effects conferred by inoculation with arbuscular mycorrhizal fungi across the host maternal genotypes at the end of the experiment (Fig. 4A & Supporting Information Table S17; *p* = 0.11, *n* = 30 maternal genotypes). Host genotypes that experienced more positive growth effects due to inoculation with mycorrhizal fungi (i.e., higher increases in leaf number in AMF relative to NM plants) experienced slightly more negative defensive outcomes (i.e., higher increases in the number of infected leaves in AMF relative to NM plants).

**Figure 4:**
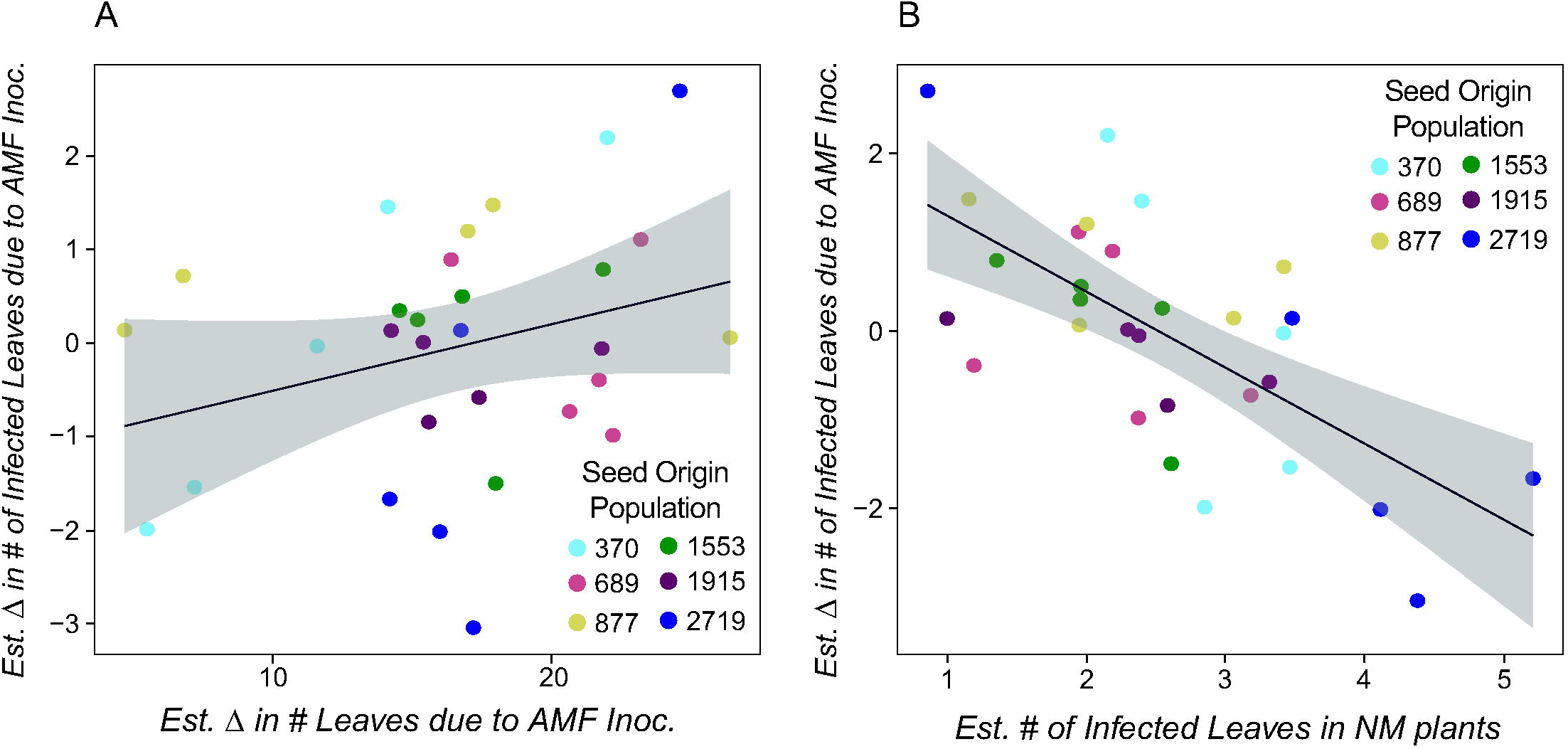
Growth and disease susceptibility are linked to defensive effects from arbuscular mycorrhizal fungi in the host genotypes. *Panel A:* In a field epidemic experiment with *Plantago lanceolata* individuals from thirty maternal genotypes, host genotypes that grew larger when inoculated with a three-species mixture of arbuscular mycorrhizal fungi experienced marginally more negative infection outcomes from exposure to *Podosphaera plantaginis* (Supporting Information Table S17; *p* = 0.11, *n* = 30 genotypes). *Panel B*: In the same experiment, host genotypes that were more susceptible to infection (when not inoculated with mycorrhizae) received stronger disease protection effects from inoculation with mycorrhizal fungi (Supporting Information Table S18; *p* < 0.001, *n* = 30 genotypes). In *A & B*, each point represents one maternal genotype, originating from one of six seed origin populations. The y-axis represents defensive effects due to mycorrhizae, i.e., the estimated change in the mean number of infected leaves in each genotype due to mycorrhizal inoculation. The shaded area represents a 95 % confidence interval. In *A*, the x-axis represents growth effects due to mycorrhizae, i.e., the estimated change in mean host size (leaf number) in each genotype due to mycorrhizal inoculation. Changes in host size were estimated before pathogen exposure in variable field conditions. In *B*, the x-axis represents host disease susceptibility, i.e., the estimated mean number of infected leaves in infected NM plants in each genotype at the peak of the epidemic: more positive x-axis values indicate more severe infections in the absence of the mutualist.

### Are disease susceptibility and defensive effects from mycorrhizal inoculation linked in host genotypes?

We found a strong relationship between host disease susceptibility and the magnitude of the defensive effect received from mycorrhizal inoculation in the maternal genotypes (Fig. 4B & Supporting Information Table S18; *p* < 0.001, *n* = 29 maternal genotypes). At the peak of the epidemic, host genotypes that suffered more severe infections (i.e., had more infected leaves) when not inoculated with mycorrhizal fungi experienced greater reductions in disease severity (i.e., in the number of infected leaves) when inoculated with mycorrhizal fungi.

## Discussion

While both plant-associated pathogenic and mutualistic microbes are ubiquitous across ecosystems, how they interact to determine patterns of infection in genetically diverse host populations is not known. In a field experiment placing wild hosts of thirty maternal genotypes in naturally occurring pathogen epidemics, we found that inoculation with a three-species mixture of arbuscular mycorrhizal fungi produced benefits and risks that influenced aboveground patterns of host infection. Mycorrhizal inoculation increased growth in hosts of nearly every genotype, but also increased infection rates from a foliar fungal pathogen. The effects of mycorrhizal inoculation on disease severity varied over the course of the epidemic, with both protective and negative effects occurring among the host genotypes. Moreover, disease susceptibility and mycorrhizae-derived defense effects appeared to be linked in the host genotypes: more susceptible host genotypes (in the absence of the mutualists) received stronger protection against disease when inoculated with mycorrhizal fungi. Arbuscular mycorrhizal fungi have been shown to protect agricultural plants against belowground (Azcón-Aguilar & Barea, 1996) and foliar (Fiorilli *et al*., 2018, Pozo de la Hoz *et al*., 2021) pathogens in controlled laboratory conditions, but our results provide the first evidence of how microbial mutualists can shift patterns of host infection in genetically diverse wild populations under pathogen attack. Mycorrhizal fungi also appeared to be linked to a growth-defense trade-off in the hosts: mycorrhizal-inoculated plants grew larger and became more infected by the pathogen, with the host genotypes that obtained the greatest growth benefit from mycorrhizal fungi also suffering the largest increases in disease. Together, our results underscore that under natural ecological and epidemic conditions mycorrhizal fungi produce a complex and temporally variable array of positive and negative effects on host growth and infection.

Inoculation with mycorrhizal fungi provided growth benefits to host plants from every population and maternal genotype. The magnitude of these effects varied among hosts from different genetic origins before exposure to the field epidemics but homogenized over time in field conditions. Due to the positive relationship between plant size and survival, growth is often intrinsically linked to host fitness (Harper & White, 1974). Our results are consistent with studies reporting evidence for the importance of mycorrhizal fungi in determining host growth and/or fitness and showing variability in such effects among host genotypes (Rasmussen *et al*., 2017, Qin *et al*., 2021). In addition, susceptibility to the pathogen epidemics in the field varied among hosts of different genetic origins, consistent with other studies in this pathosystem (Laine, 2005, Laine, 2007a, Tack *et al*., 2014, Susi & Laine, 2015) and the broader wild plant disease literature (Carlsson-Granér, 1997, Price *et al*., 2004). Together, these results provide evidence for the importance of genotype in mediating host-microbe interactions (Eck *et al*. 2019, Sallinen *et al*., 2020), although more generalist interactions can also certainly occur (Gilbert & Webb, 2007, Halbritter *et al*., 2012, Hersh *et al*., 2012).

Building upon such studies, we show evidence of changes in host-parasite interactions due to prior inoculation with microbial mutualists. Inoculation experiments with agricultural plant species and their pathogens have shown that arbuscular mycorrhizal fungi can reduce disease incidence or severity (Norman *et al*., 1996; Pozo *et al*., 2002; Li *et al*., 2010; Song *et al*., 2015; Berdeni *et al*., 2018), However, symbiotic relationships in cultivars may differ from those of wild plants (Xing *et al*., 2012); thus, it is not straightforward to predict responses in wild populations from controlled agricultural trials. In this study, mycorrhizal inoculation increased infection rates in hosts of a wild plant species and produced variable effects on disease severity in hosts of different genotypes and over the course of the epidemic. By the end of the experiment, mycorrhizal inoculation was weakly linked to higher numbers of infected leaves in diseased hosts (though variation among genotypes remained). Our results suggest that environmental, temporal, and genetic contexts may alter the potential defensive effects related to mycorrhizal association and are consistent with other experimental studies showing that mycorrhizal effects on host infection may vary among host genotypes (Mark & Cassells 1996; Steinkellner *et al*. 2012). However, in our experiment the effects of mycorrhizal inoculation on infection were weaker than the effects on host growth. Additional studies where mycorrhizal colonization is explicitly quantified are needed to more reliably assess the contribution of the symbiont to host growth, infection, and fitness. The potential risks of mycorrhizal association in wild plants are relevant for theoretical studies which speculate as to how mycorrhizal fungi might influence host fitness in the presence of pathogens and affect plant population and community dynamics (Bachelot *et al*., 2015).

In many free-living organisms, an evolutionary trade-off exists between growth and defense (Herms & Mattson, 1992). In our experiment, additional insights can be gained by examining whether changes in host growth due to mutualism are related to changes in host defense across genotypes. In our experiment, mycorrhizal-inoculated hosts consistently grew larger and were more likely to become infected by the pathogen. This could occur if increases in host size increase pathogen encounter rates, as could be expected for pathogen species with wind-or passively-dispersed spores (like *Po. plantaginis*). In addition, host genotypes that experienced larger growth benefits from mycorrhizal inoculation suffered marginally larger increases in disease severity (though there was considerable variation in this effect). That some host genotypes grew larger and experienced reductions in disease severity following mycorrhizal inoculation suggests that defense priming could occur occasionally. Infected AMF plants also had lower proportions of infected leaves than NM plants (relative to their size), though it is unclear whether this may offset the costs of having higher numbers of infected leaves in this pathosystem. Changes in host tolerance to pathogens following mycorrhizal inoculation could also explain increases in infected leaf numbers, though mycorrhizal fungi are generally expected to improve host nutritional status or increase leaf toughness, making foliar pathogen spread more difficult (Meier & Hunter, 2018). Thus, it is likely that differences in host infection between AMF and NM plants in our experiment were mediated by increases in host size (as host size also predicted some aspects of infection) or defense priming in some genotypes.

In addition, our findings indicate that arbuscular mycorrhizal fungi could help susceptible host genotypes compensate for lack of innate resistance while placing costs on well-defended genotypes. We found that the magnitude of defense effects following mycorrhizal inoculation were linked to pathogen susceptibility in the host genotypes: host genotypes that were susceptible to more severe infections received stronger disease protection effects when inoculated with the mutualist. In contrast, more resistant host genotypes were likely to slightly increase in disease load when inoculated. Thus, inoculation with mycorrhizal fungi tended to equalize disease severity between more resistant and more susceptible host genotypes, potentially reducing the relative importance of host genetic resistance in determining pathogen effects. However, the linkages between disease susceptibility and mycorrhizal defense effects in our study (as well as between host growth and these effects) were revealed post-hoc and should be confirmed with experiments designed to test these hypotheses. If confirmed, it could indicate that host genotypes may experience trade-offs in investment in genetic resistance vs. mycorrhizal-mediated resistance. It could also suggest that mycorrhizal association may have been selected for and maintained in host populations partially because it increases the fitness of susceptible host genotypes. In this way, mycorrhizal protective effects could contribute to the maintenance of diversity in host genetic resistance within and among populations (Laine, 2004; Laine 2007a, Jousimo *et al*. 2014).

Variation among host populations and genotypes in mycorrhizal benefits and risks could be due to several factors. These factors include intraspecific differences in hosts’ ability to form associations with and derive function from different mycorrhizal species, differences in mycorrhizal colonization rates or community composition, environmental variation, or differences in host-pathogen population dynamics over time. Prior studies demonstrate variation in arbuscular mycorrhizal colonization rates among individuals within species – variation which is thought to have a partially genetic component (Plouznikoff *et al*. 2019; Pawlowski *et al*. 2020). Though we observed clear differences in host growth and infection as a result of the mycorrhizal inoculation treatment, data on mycorrhizal colonization are needed to confirm mycorrhizae as the mechanism underlying the observed effects. Environmental conditions may also impact plant-microbe interactions (Santoyo *et al*., 2017). Consistent with other studies in this pathosystem, infection rates varied among field populations and changed over time (Eck *et al*., 2022). In contrast, the effects of mycorrhizal inoculation on hosts were similar among field populations, though changes over time also occurred. Growth benefits following inoculation with mycorrhizal fungi varied among host populations and genotypes before pathogen exposure but became homogenous over time in the field conditions. Changes in host infection rate and disease severity due to mycorrhizal inoculation were also similar among field populations but increased over time. Together, these results suggest that mycorrhizal effects on hosts are more temporally than environmentally sensitive. There is also some chance that environmental mycorrhizal spores could have come in to contact with our experimental soils, such that true levels of mycorrhizal colonization in the non-mycorrhizal treatment could be low or some (rather than none), and the mycorrhizal communities in the experimental pots could contain species that were not inoculated. The limited duration of the field experiment reduces the likelihood that this could cause strong effects (Sanders & Sheikh, 1986); however, if it occurred, it should have occurred evenly among treatments and seed sources. Future studies in which mycorrhizal composition, colonization rates, and function are quantified in genetically diverse hosts and in variable environmental conditions over time are necessary to corroborate this work and disentangle the effect of mycorrhizal growth and defensive effects from intraspecific variation in host growth and resistance.

Together, our results suggest that symbiosis with arbuscular mycorrhizal fungi produces benefits and alters infection risks from a pathogen during natural epidemics in genetically variable host populations. Altered patterns of host growth and infection due to mutualist species may cascade to affect patterns of abundance, diversity, and distribution of the associated organisms, as well as ecosystem processes (Brown *et al*., 2001). Moreover, we are beginning to acknowledge the importance of direct and indirect microbial interactions within hosts in determining host fitness and parasite population dynamics (Kemen, 2014; Kemen *et al*., 2015; Kroll *et al*., 2017). Belowground soil and rhizosphere processes may also affect aboveground interactions with pathogens and herbivores, and vice versa (van der Putten *et al*., 2001; Wardle *et al*., 2004; Frew, 2021), and mutualists and parasites may act simultaneously to determine host fitness (Bezemer & van Dam, 2005). Our results also highlight the importance of characterizing host-microbe interactions under natural conditions and temporal interaction sequences, which will allow more precise modeling of epidemiological and ecological community dynamics. In addition, growth and defensive effects due to mutualism should ultimately affect the co-evolutionary trajectories of the associated organisms, because we expect organisms to be adapted to events that are common over evolutionary time spans (van Dam & Heil, 2011). Understanding how mutualism alters host susceptibility and parasite interactions will be important for understanding, predicting, and managing disease in ecological communities and in agriculture.

## Acknowledgements

We would like to thank the University of Oulu Botanical Gardens for providing glasshouse facilities, Riitta Ovaska for assistance during the field experiment, Dr. Mauritz Vestberg for the fungal isolates, and the anonymous reviewers for providing constructive feedback. This work was funded by grants from the European Research Council (Consolidator Grant RESISTANCE: LS8 & ERC-2016-COG) and the Academy of Finland (MULTA #327222 & #296686) to A-LL, and the Forschungskredit of the University of Zurich (FK-20-108 & FK-20-118) to JLE.

## Author Contributions

A-LL and M-MK designed and conducted the experiment. JLE and A-LL analyzed the data. JLE wrote the first draft of the manuscript. All authors contributed to and approved the final version.

## Data Availability

The data that supports the findings of this study are available in the supplementary material of this article

## Legend of Supporting Information

Dataset S1 Growth and disease data from the field epidemic experiment.

**Fig. S1** Photograph of *Pl. lanceolata* showing signs of infection with *Po. plantaginis*.

**Fig. S2** Growth of AMF plants relative to NM plants in each seed origin population at the beginning of the field epidemic experiment.

**Fig. S3** Biomass was increased in AMF plants relative to NM plants at the end of the field epidemic experiment.

**Table S1** Effects of mycorrhizal inoculation and host genetic origin on the growth of healthy individuals at the start of the field epidemic experiment.

**Table S2** Factors influencing the growth of six seed origin populations at the end of the field epidemic experiment.

**Table S3** Factors influencing the growth of thirty maternal genotypes at the end of the field epidemic experiment.

**Table S4** Effect of mycorrhizal association on final harvest biomass in a subset of the experimental individuals.

**Table S5** Factors influencing host infection rate in six seed origin populations during the peak of the field epidemic experiment.

**Table S6** Factors influencing host infection rate in thirty maternal genotypes during the peak of the field epidemic experiment.

**Table S7** Factors influencing host infection rate in six seed origin populations at the end of the field epidemic experiment.

**Table S8** Factors influencing host infection rate in thirty maternal genotypes at the end of the field epidemic experiment.

**Table S9** Factors influencing the proportion of infected leaves in hosts in six seed origin populations during the epidemic peak.

**Table S10** Factors influencing the proportion of infected leaves in hosts in thirty maternal genotypes at the epidemic peak.

**Table S11** Factors influencing the proportion of infected leaves in hosts in six seed origin populations at the end of the field epidemic experiment.

**Table S12** Factors influencing the proportion of infected leaves in hosts in thirty maternal genotypes at the end of the field epidemic experiment.

**Table S13** Factors influencing the number of infected leaves in hosts in six seed origin populations during the epidemic peak.

**Table S14** Factors influencing the number of infected leaves in hosts in thirty maternal genotypes at the peak of the field epidemic experiment.

**Table S15** Factors influencing the number of infected leaves in hosts in six seed origin populations at the end of the field epidemic experiment.

**Table S16** Factors influencing the number of infected leaves in hosts in thirty maternal genotypes at the end of the field epidemic experiment.

**Table S17** Relationship between host growth and defensive effects due to mycorrhizal inoculation in the maternal genotypes.

**Table S18** Relationship between disease susceptibility and defensive effects due to mycorrhizal inoculation in the maternal genotypes.

