## Supporting Information for "Arbuscular mycorrhizal fungi influence host infection during epidemics in a wild plant pathosystem"

New Phytologist

***New Phytologist* Supporting Information**

Article acceptance date: 12 July 2022

The following Supporting Information is available for this article:

**Dataset S1** Growth and disease data from the field epidemic experiment.

**Fig. S1** Photograph of *Pl. lanceolata* showing signs of infection with *Po. plantaginis*.

**Fig. S2** Growth of AMF plants relative to NM plants in each seed origin population at the beginning of the field epidemic experiment.

**Fig. S3** Biomass was increased in AMF plants relative to NM plants at the end of the field epidemic experiment.

**Table S1** Effects of mycorrhizal inoculation and host genetic origin on the growth of healthy individuals at the start of the field epidemic experiment.

**Table S2** Factors influencing the growth of six seed origin populations at the end of the field epidemic experiment.

**Table S3** Factors influencing the growth of thirty maternal genotypes at the end of the field epidemic experiment.

**Table S4** Effect of mycorrhizal association on final harvest biomass in a subset of the experimental individuals.

**Table S5** Factors influencing host infection rate in six seed origin populations during the peak of the field epidemic experiment.

**Table S6** Factors influencing host infection rate in thirty maternal genotypes during the peak of the field epidemic experiment.

**Table S7** Factors influencing host infection rate in six seed origin populations at the end of the field epidemic experiment.

**Table S8** Factors influencing host infection rate in thirty maternal genotypes at the end of the field epidemic experiment.

**Table S9** Factors influencing the proportion of infected leaves in hosts in six seed origin populations during the epidemic peak.

**Table S10** Factors influencing the proportion of infected leaves in hosts in thirty maternal genotypes at the epidemic peak.

**Table S11** Factors influencing the proportion of infected leaves in hosts in six seed origin populations at the end of the field epidemic experiment.

**Table S12** Factors influencing the proportion of infected leaves in hosts in thirty maternal genotypes at the end of the field epidemic experiment.

**Table S13** Factors influencing the number of infected leaves in hosts in six seed origin populations during the epidemic peak.

**Table S14** Factors influencing the number of infected leaves in hosts in thirty maternal genotypes at the peak of the field epidemic experiment.

**Table S15** Factors influencing the number of infected leaves in hosts in six seed origin populations at the end of the field epidemic experiment.

**Table S16** Factors influencing the number of infected leaves in hosts in thirty maternal genotypes at the end of the field epidemic experiment.

**Table S17** Relationship between host growth and defensive effects due to mycorrhizal inoculation in the maternal genotypes.

**Table S18** Relationship between disease susceptibility and defensive effects due to mycorrhizal inoculation in the maternal genotypes.

New Phytologist

**Fig. S1** Photograph of *Pl. lanceolata* showing signs of infection with *Po. plantaginis*. Infection by *Po. plantaginis* on *Pl. lanceolata* produces white, powdery spot(s) and/or blotches on the foliar tissue. Spots often grow larger over time and are visible to the naked eye. This sign of disease is characteristic of *Po. plantaginis* in the *Pl. lanceolata* populations in the Åland Islands, Finland.

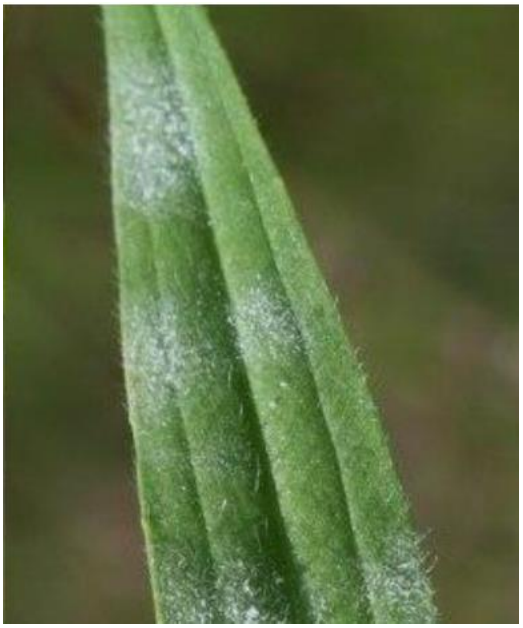

**Fig. S2** Growth of AMF plants relative to NM plants in each seed origin population at the beginning of the field epidemic experiment. Inoculation with a mixture of three arbuscular mycorrhizal fungal species increased the growth of experimental *Pl. lanceolata* individuals from six seed origin populations, relative to NM individuals, at the time of placement in the field epidemic experiment (i.e., before pathogen exposure) (Table S1; MYC: $p < 0.001$, $n = 287$ plants). Growth also varied among seed origin populations (Table S1; POP: $p < 0.001$, $n = 287$ plants). Growth was modeled as the number of leaves on each individual during the first survey of the experiment. Box notches represent a 95 % confidence interval for comparing medians. Box hinges correspond to the 1st and 3rd quartiles. Box whiskers extend to the largest and smallest value no further than 1.5 × the interquartile range from the hinges.

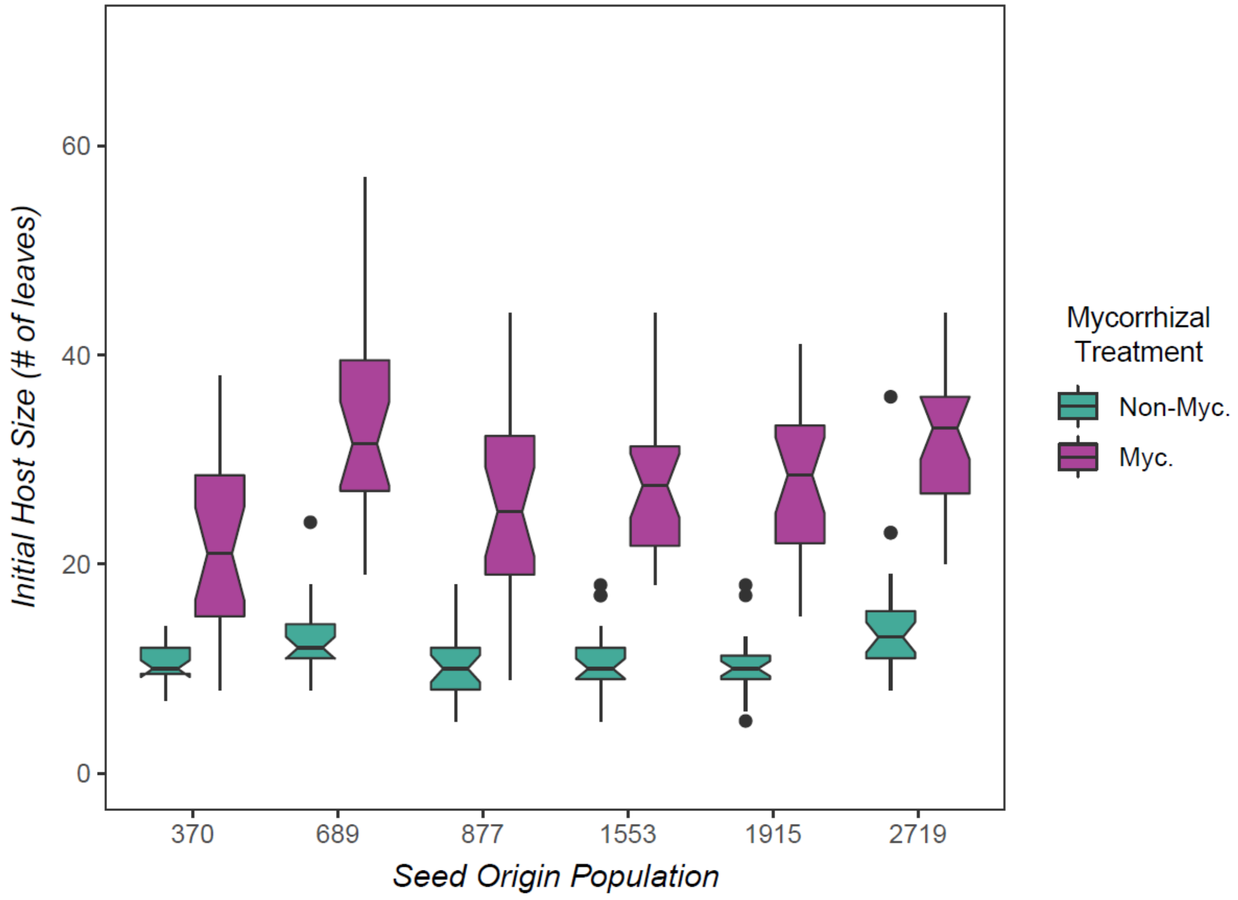

**Fig. S3** Biomass was increased in AMF plants relative to NM plants at the end of the field epidemic experiment. Inoculation with a mixture of three arbuscular mycorrhizal fungal species increased the dry aboveground biomass of *Pl. lanceolata* individuals, relative to NM plants, at the end of the natural epidemic field experiment (i.e., post-pathogen exposure) (Table S4; MYC: $W = 11$, $p < 0.001$, $n = 41$ plants). Biomass was measured on a haphazardly selected subset of 21 AMF plants and 20 NM plants. Box hinges correspond to the 1st and 3rd quartiles. Box whiskers extend to the largest and smallest value no further than 1.5 × the interquartile range from the hinges.

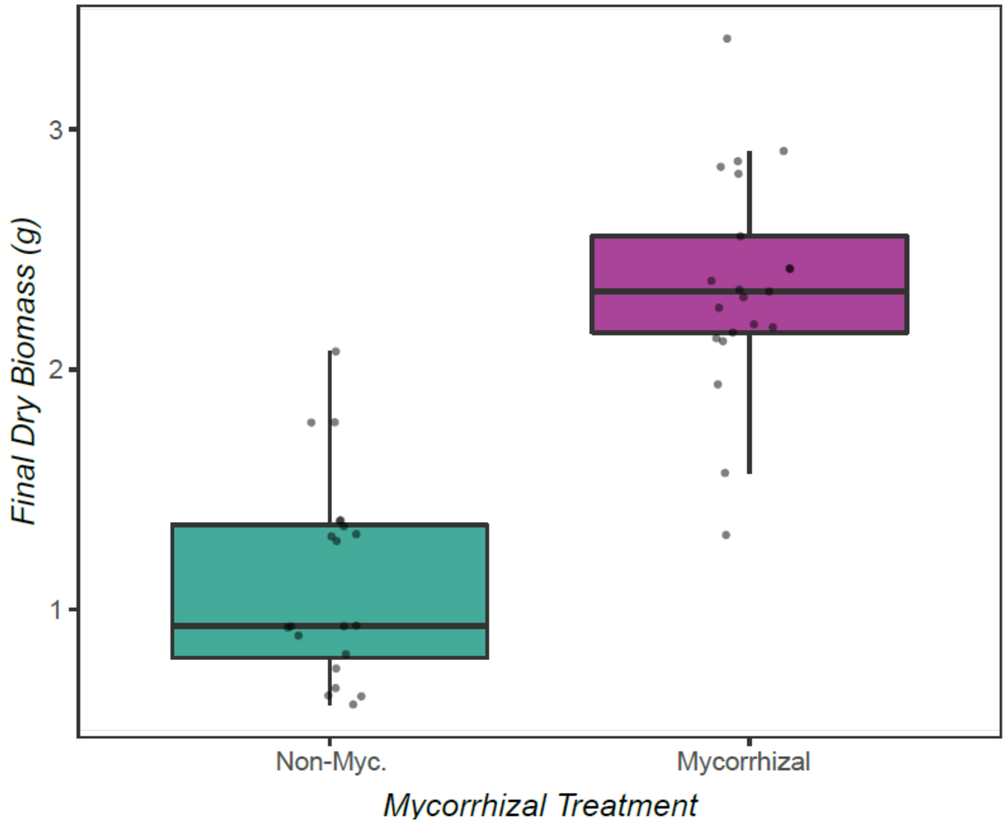

**Table S1** Effects of mycorrhizal inoculation and host genetic origin on the growth of healthy individuals at the start of the field epidemic experiment. We examined the effects of inoculation with arbuscular mycorrhizal fungi, seed origin population (Table S1A) or maternal genotype (Table S1B) and their interaction on the growth of healthy experimental individuals using a negative binomial generalized linear model ($n$ = 287 individuals). Growth was measured as the number of leaves on each individual before pathogen exposure (i.e., at the start of the field epidemic experiment). Non-significant interaction terms were removed from the final model. Analysis of deviance results are shown in the table below, including degrees of freedom (d.f.), deviance, residual d.f., residual deviance, $F$-score, and $p$-value for each factor. Factors for which $p < 0.05$ are highlighted in bold.

**Table S1A: Seed Origin Population**

| *Analysis of Deviance* | *d.f.* | *Deviance* | *Residual d.f.* | *Residual Deviance* | *F* | *p* |
|---|---|---|---|---|---|---|
| *Null* | - | - | 286 | 932.20 | - | - |
| Mycorrhizal Inoculation | 1 | 605.60 | 285 | 326.60 | 605.60 | **$< 2.2 \times e^{-16}$** |
| Seed Origin Population | 5 | 57.29 | 280 | 269.32 | 11.46 | **$4.42 \times e^{-11}$** |

**Table S1B: Maternal Genotype**

| | *d.f.* | *Deviance* | *Residual d.f.* | *Residual Deviance* | *F* | *p* |
|---|---|---|---|---|---|---|
| *Null* | - | - | 286 | 1249.29 | - | - |
| Mycorrhizal Inoculation | 1 | 806.05 | 285 | 443.24 | 806.05 | **$< 2.2 \times e^{-16}$** |
| Maternal Genotype | 29 | 132.72 | 256 | 310.50 | 4.58 | **$3.21 \times e^{-15}$** |
| Myc. Inoc. × Mat. Gen. | 29 | 56.23 | 227 | 254.27 | 1.94 | **$1.77 \times e^{-03}$** |

**Table S2** Factors influencing the growth of six seed origin populations at the end of the field epidemic experiment. We examined the effects of inoculation with arbuscular mycorrhizal fungi, seed origin population, field epidemic population, host initial size (Table S2B), and their interactions on the growth of experimental individuals at the end of the field epidemic experiment (i.e., after pathogen exposure) using a negative binomial generalized linear model ($n$ = 287 individuals). Host growth was modeled as the number of leaves on each individual during the final survey of the experiment. Table S2B shows the results of the model with host initial size included. Non-significant interaction terms were removed from the final model. Analysis of deviance results are shown in the table below, including degrees of freedom (d.f.), deviance, residual d.f., residual deviance, $F$-score, and $p$-value for each factor. Factors for which $p < 0.05$ are highlighted in bold.

### Table S2A: Without Initial Size

| *Analysis of Deviance* | *d.f.* | *Deviance* | *Residual d.f.* | *Residual Deviance* | *F* | *p* |
|---|---|---|---|---|---|---|
| *Null* | - | - | 286 | 722.89 | - | - |
| Mycorrhizal Inoculation | 1 | 433.81 | 285 | 289.08 | 433.81 | **< 2.2 x e$^{-16}$** |
| Seed Origin Population | 5 | 23.25 | 280 | 265.82 | 4.65 | **3.02 x e$^{-04}$** |
| Field Epidemic Population | 2 | 1.60 | 278 | 265.23 | 0.78 | 0.45 |

### Table S2B: Including Initial Size

| *Analysis of Deviance* | *d.f.* | *Deviance* | *Residual d.f.* | *Residual Deviance* | *F* | *p* |
|---|---|---|---|---|---|---|
| *Null* | - | - | 286 | 1322.82 | - | - |
| Mycorrhizal Inoculation | 1 | 771.61 | 285 | 551.21 | 771.61 | **< 2.2 x e$^{-16}$** |
| Host Initial Size | 1 | 295.70 | 284 | 255.51 | 295.70 | **< 2.2 x e$^{-16}$** |
| Seed Origin Population | 5 | 3.81 | 279 | 251.71 | 0.76 | 0.58 |
| Field Epidemic Population | 2 | 3.17 | 277 | 248.53 | 1.59 | 0.20 |
| Myc. Inoc. × Host Init. Size | 1 | 13.58 | 276 | 234.95 | 13.58 | **2.28 x e$^{-04}$** |

New Phytologist

**Table S3** Factors influencing the growth of thirty maternal genotypes at the end of the field epidemic experiment. We examined the effects of inoculation with arbuscular mycorrhizal fungi, maternal genotype, field epidemic population, and a three-way interaction between all factors on the growth of experimental individuals at the end of the field epidemic experiment (i.e., after pathogen exposure) using a negative binomial generalized linear model ($n$ = 287 individuals). Host growth was modeled as the number of leaves on each individual during the final survey of the experiment. A model including host initial size was not constructed due to problems concerning model fit. Non-significant interaction terms were removed from the final model. Analysis of deviance results are shown in the table below, including degrees of freedom (d.f.), deviance, residual d.f., residual deviance, $F$-score, and $p$-value for each factor. Factors for which $p < 0.05$ are highlighted in bold.

| ***Analysis of Deviance*** | ***d.f.*** | ***Deviance*** | ***Residual d.f.*** | ***Residual Deviance*** | ***F*** | ***p*** |
|---|---|---|---|---|---|---|
| *Null* | - | - | 286 | 820.39 | - | - |
| Mycorrhizal Inoculation | 1 | 490.67 | 285 | 329.71 | 490.67 | **< 2.2 x $e^{-16}$** |
| Maternal Genotype | 29 | 62.79 | 256 | 266.93 | 2.17 | **2.75 x $e^{-04}$** |
| Field Epidemic Population | 2 | 2.90 | 254 | 264.03 | 1.45 | 0.23 |

**Table S4** Effect of mycorrhizal association on final harvest biomass in a subset of the experimental individuals. We examined the effect of the mycorrhizal inoculation treatment on the final dry aboveground biomass of experimental individuals at the end of the field epidemic experiment (i.e., post-pathogen exposure) using a Mann-Whitney U test ($n$ = 41 individuals: 21 AMF and 20 NM). The Wilcoxon $W$-statistic and $p$-value resulting from this test are reported in the table below.

| ***Mann-Whitney U Test*** | ***W*** | ***p*** |
|---|---|---|
| Mycorrhizal Inoculation | 11 | **1.45 x e$^{-09}$** |

**Table S5** Factors influencing host infection rate in six seed origin populations during the peak of the field epidemic experiment. We examined the effects of inoculation with arbuscular mycorrhizal fungi, seed origin population, field epidemic population, host size (Table S5B) and their interactions on the infection status of *Pl. lanceolata* individuals during the peak epidemic period in the field experiment using a generalized logistic regression model ($n$ = 287 individuals). Table S5B shows the results of the model with host size included. Non-significant interaction terms were removed from the final models. Analysis of deviance results are shown in the table below, including degrees of freedom (d.f.), deviance, residual d.f., residual deviance, $F$-score, and $p$-values. Effects for which $p < 0.05$ are highlighted in bold.

**Table S5A: Without Host Size**

| **Analysis of Deviance** | ***d.f.*** | ***Deviance*** | ***Resid. d.f.*** | ***Resid. Dev.*** | ***F*** | ***p*** |
|---|---|---|---|---|---|---|
| Null Model | - | - | 286 | 333.81 | - | - |
| Mycorrhizal Inoculation | 1 | 2.89 | 285 | 330.92 | 2.85 | 0.09 |
| Seed Origin Population | 5 | 4.90 | 280 | 326.02 | 0.97 | 0.44 |
| Field Epidemic Population | 2 | 7.57 | 278 | 318.45 | 3.74 | **0.03** |

**Table S5B: With Host Size**

| **Analysis of Deviance** | ***d.f.*** | ***Deviance*** | ***Resid. d.f.*** | ***Resid. Dev.*** | ***F*** | ***p*** |
|---|---|---|---|---|---|---|
| Null Model | - | - | 286 | 331.81 | - | - |
| Mycorrhizal Inoculation | 1 | 2.89 | 285 | 330.92 | 2.80 | 0.10 |
| Seed Origin Population | 5 | 4.90 | 280 | 326.02 | 0.95 | 0.45 |
| Field Epidemic Population | 2 | 7.57 | 278 | 318.45 | 3.68 | **0.03** |
| Host Size | 1 | 0.48 | 277 | 317.97 | 0.47 | 0.50 |

**Table S6** Factors influencing host infection rate in thirty maternal genotypes during the peak of the field epidemic experiment. We examined the effects of inoculation with arbuscular mycorrhizal fungi, maternal genotype, field epidemic population, host size (Table S6B) and their interactions on the infection status of *Pl. lanceolata* individuals during the peak epidemic period in the field experiment using a generalized logistic regression model ($n$ = 287 individuals). Table S6B shows the results of the model with host size included. Non-significant interaction terms were removed from the final model. Analysis of deviance results are shown in the table below, including degrees of freedom (d.f.), deviance, residual d.f., residual deviance, $F$-score, and $p$-values. Effects for which $p < 0.05$ are highlighted in bold.

#### Table S6A: Without Host Size

| Analysis of Deviance | *d.f.* | *Deviance* | *Resid. d.f.* | *Resid. Dev.* | *F* | *p* |
|---|---|---|---|---|---|---|
| Null Model | - | - | 286 | 333.81 | - | - |
| Mycorrhizal Inoculation | 1 | 2.89 | 285 | 330.92 | 2.56 | 0.11 |
| Maternal Genotype | 29 | 35.09 | 256 | 295.84 | 1.07 | 0.37 |
| Field Epidemic Population | 2 | 9.90 | 254 | 285.94 | 4.39 | **0.01** |

#### Table S6B: With Host Size

| Analysis of Deviance | *d.f.* | *Deviance* | *Resid. d.f.* | *Resid. Dev.* | *F* | *p* |
|---|---|---|---|---|---|---|
| Null Model | - | - | 286 | 333.81 | - | - |
| Mycorrhizal Inoculation | 1 | 2.89 | 285 | 330.92 | 2.55 | 0.11 |
| Maternal Genotype | 29 | 35.09 | 256 | 295.84 | 1.07 | 0.37 |
| Field Epidemic Population | 2 | 9.90 | 254 | 285.94 | 4.39 | **0.01** |
| Host Size | 1 | 0.04 | 253 | 285.90 | 0.04 | 0.85 |

**Table S7** Factors influencing host infection rate in six seed origin populations at the end of the field epidemic experiment. We examined the effects of inoculation with arbuscular mycorrhizal fungi, seed origin population, field epidemic population, host size (Table S7B) and their interactions on the infection status of *Pl. lanceolata* individuals at the end of the field epidemic experiment using a generalized logistic regression model ($n$ = 286 individuals). Table S7B shows the results of the model with host initial size included. Non-significant interaction terms were removed from the final model. Analysis of deviance results are shown in the table below, including degrees of freedom (d.f.), deviance, residual d.f., residual deviance, $F$-score, and $p$-values. Effects for which $p < 0.05$ are highlighted in bold.

**Table S7A: Without Host Size**

| **Analysis of Deviance** | ***d.f.*** | ***Deviance*** | ***Resid. d.f.*** | ***Resid. Dev.*** | ***F*** | ***p*** |
|---|---|---|---|---|---|---|
| Null Model | - | - | 285 | 367.67 | - | - |
| Mycorrhizal Inoculation | 1 | 4.34 | 284 | 363.34 | 4.16 | **0.04** |
| Seed Origin Population | 5 | 4.61 | 279 | 358.73 | 0.88 | 0.49 |
| Field Epidemic Population | 2 | 46.91 | 277 | 311.82 | 22.50 | **$8.83 \times e^{-10}$** |

**Table S7B: With Host Size**

| **Analysis of Deviance** | ***d.f.*** | ***Deviance*** | ***Resid. d.f.*** | ***Resid. Dev.*** | ***F*** | ***p*** |
|---|---|---|---|---|---|---|
| Null Model | - | - | 285 | 367.67 | - | - |
| Mycorrhizal Inoculation | 1 | 4.34 | 284 | 363.34 | 4.07 | **0.04** |
| Seed Origin Population | 5 | 4.61 | 279 | 358.73 | 0.86 | 0.51 |
| Field Epidemic Population | 2 | 46.91 | 277 | 311.82 | 22.00 | **$1.36 \times e^{-09}$** |
| Host Size | 1 | 3.93 | 276 | 307.89 | 3.69 | 0.06 |

**Table S8** Factors influencing host infection rate in thirty maternal genotypes at the end of the field epidemic experiment. We examined the effects of inoculation with arbuscular mycorrhizal fungi, maternal genotype, field epidemic population, host size (Table S8B), and their interactions on the infection status of *Pl. lanceolata* individuals at the end of the field epidemic experiment using a generalized logistic regression model ($n$ = 286 individuals). Table S8B shows the results of the model with host initial size included. Non-significant interaction terms were removed from the final model. Analysis of deviance results are shown in the table below, including degrees of freedom (d.f.), deviance, residual d.f., residual deviance, $F$-score, and $p$-values. Effects for which $p < 0.05$ are highlighted in bold.

**Table S8A: Without Host Size**

| **Analysis of Deviance** | ***d.f.*** | ***Deviance*** | ***Resid. d.f.*** | ***Resid. Dev.*** | ***F*** | ***p*** |
|---|---|---|---|---|---|---|
| Null Model | - | - | 285 | 367.67 | - | - |
| Mycorrhizal Inoculation | 1 | 4.34 | 284 | 363.34 | 3.93 | **0.05** |
| Maternal Genotype | 29 | 34.81 | 255 | 328.53 | 1.09 | 0.35 |
| Field Epidemic Population | 2 | 51.43 | 253 | 277.10 | 23.32 | **5.07 x $e^{-10}$** |

**Table S8B: With Host Size**

| **Analysis of Deviance** | ***d.f.*** | ***Deviance*** | ***Resid. d.f.*** | ***Resid. Dev.*** | ***F*** | ***p*** |
|---|---|---|---|---|---|---|
| Null Model | - | - | 285 | 367.67 | - | - |
| Mycorrhizal Inoculation | 1 | 4.34 | 284 | 363.34 | 4.01 | **0.05** |
| Maternal Genotype | 29 | 34.81 | 255 | 328.53 | 1.11 | 0.33 |
| Field Epidemic Population | 2 | 51.43 | 253 | 277.10 | 23.78 | **3.47 x $e^{-10}$** |
| Host Size | 1 | 4.68 | 252 | 272.42 | 4.33 | **0.04** |

**Table S9** Factors influencing the proportion of infected leaves in hosts in six seed origin populations during the epidemic peak. We examined the effects of inoculation with arbuscular mycorrhizal fungi, seed origin population, field epidemic population, host size (Table S9B), and their interactions on the proportion of infected leaves in Pl. lanceolata individuals during the peak of the field epidemic experiment using a generalized logistic regression model ($n$ = 210 individuals). Table S9B shows the results of the model with host initial size included. Non-significant interaction terms were removed from the final model. Analysis of deviance results are shown in the table below, including degrees of freedom (d.f.), deviance, residual d.f., residual deviance, $F$-score, and $p$-values. Effects for which $p < 0.05$ are highlighted in bold.

#### Table S9A: Without Host Size

| Analysis of Deviance | *d.f.* | *Deviance* | *Resid. d.f.* | *Resid. Dev.* | *F* | *p* |
|---|---|---|---|---|---|---|
| Null Model | - | - | 209 | 334.85 | - | - |
| Mycorrhizal Inoculation | 1 | 107.15 | 208 | 227.70 | 107.15 | **< 2.2 x $e^{-16}$** |
| Field Epidemic Population | 2 | 5.66 | 206 | 222.04 | 2.83 | 0.06 |
| Seed Origin Population | 5 | 17.58 | 201 | 204.46 | 3.52 | **0.004** |
| Myc. Inoc. × Field Pop. | 2 | 5.66 | 199 | 198.80 | 2.83 | 0.06 |

#### Table S9B: With Host Size

| Analysis of Deviance | *d.f.* | *Deviance* | *Resid. d.f.* | *Resid. Dev.* | *F* | *p* |
|---|---|---|---|---|---|---|
| Null Model | - | - | 209 | 334.85 | - | - |
| Mycorrhizal Inoculation | 1 | 107.15 | 208 | 227.7 | 107.15 | **< 2.2 x $e^{-16}$** |
| Field Epidemic Population | 2 | 5.66 | 206 | 222.04 | 2.83 | 0.06 |
| Seed Origin Population | 5 | 17.58 | 201 | 204.46 | 3.52 | **0.004** |
| Host Size | 1 | 19.57 | 200 | 184.89 | 19.57 | **9.70 x $e^{-06}$** |
| Myc. Inoc. × Field Pop. | 2 | 5.05 | 198 | 179.84 | 2.53 | 0.08 |

**Table S10** Factors influencing the proportion of infected leaves in hosts in thirty maternal genotypes at the epidemic peak. We examined the effects of inoculation with arbuscular mycorrhizal fungi, maternal genotype, field epidemic population, host size (Table S10B), and their interactions on the proportion of infected leaves in *Pl. lanceolata* individuals during the peak of the field epidemic experiment using a generalized logistic regression model ($n = 210$ individuals). Table S10B shows the results of the model with host initial size included. Non-significant interaction terms were removed from the final model. Analysis of deviance results are shown in the table below, including degrees of freedom (d.f.), deviance, residual d.f., residual deviance, $F$-score, and $p$-values. Effects for which $p < 0.05$ are highlighted in bold.

**Table S10A: Without Host Size**

| Analysis of Deviance | *d.f.* | *Deviance* | *Resid. d.f.* | *Resid. Dev.* | *F* | *p* |
|---|---|---|---|---|---|---|
| Null Model | - | - | 209 | 334.85 | - | - |
| Mycorrhizal Inoculation | 1 | 107.154 | 208 | 227.70 | 107.15 | **$< 2.2 \times e^{-16}$** |
| Field Epidemic Population | 2 | 5.66 | 206 | 222.04 | 2.83 | 0.06 |
| Maternal Genotype | 29 | 46.82 | 177 | 175.22 | 1.61 | **0.02** |
| Field Pop. × Mat. Gen. | 56 | 76.34 | 121 | 98.88 | 1.36 | **0.04** |
| Myc Inoc. × Field Pop. | 2 | 9.44 | 119 | 89.44 | 4.72 | **0.009** |

**Table S10B: With Host Size**

| Analysis of Deviance | *d.f.* | *Deviance* | *Resid. d.f.* | *Resid. Dev.* | *F* | *p* |
|---|---|---|---|---|---|---|
| Null Model | - | - | 209 | 334.85 | - | - |
| Mycorrhizal Inoculation | 1 | 107.15 | 208 | 227.70 | 107.15 | **$< 2.2 \times e^{-16}$** |
| Field Epidemic Population | 2 | 5.66 | 206 | 222.04 | 2.83 | 0.06 |
| Maternal Genotype | 29 | 46.82 | 177 | 175.22 | 1.61 | **0.02** |
| Host Size | 1 | 15.58 | 176 | 159.64 | 15.58 | **$7.92 \times e^{-05}$** |
| Myc Inoc. × Field Pop. | 2 | 4.81 | 174 | 154.83 | 2.40 | 0.09 |
| Field Pop. × Mat. Gen. | 56 | 71.82 | 118 | 83.02 | 1.28 | 0.08 |

**Table S11** Factors influencing the proportion of infected leaves in hosts in six seed origin populations at the end of the field epidemic experiment. We examined the effects of inoculation with arbuscular mycorrhizal fungi, seed origin population, field epidemic population, host size (Table S11B), and their interactions on the proportion of infected leaves in *Pl. lanceolata* individuals at the end of the field epidemic experiment using a generalized logistic regression model ($n$ = 98 individuals). Non-significant interaction terms were removed from the final model. Table S11B shows the results of the model with host initial size included. Analysis of deviance results are shown in the table below, including degrees of freedom (d.f.), deviance, residual d.f., residual deviance, $F$-score, and $p$-values. Effects for which $p < 0.05$ are highlighted in bold.

**Table S11A: Without Host Size**

| Analysis of Deviance | *d.f.* | *Deviance* | *Resid. d.f.* | *Resid. Dev.* | *F* | *p* |
|---|---|---|---|---|---|---|
| Null Model | - | - | 97 | 83.94 | - | - |
| Mycorrhizal Inoculation | 1 | 9.16 | 96 | 74.78 | 9.16 | **0.003** |
| Field Epidemic Population | 2 | 3.08 | 94 | 71.70 | 1.54 | 0.21 |
| Seed Origin Population | 5 | 6.94 | 89 | 64.76 | 1.39 | 0.22 |

**Table S11B: With Host Size**

| Analysis of Deviance | *d.f.* | *Deviance* | *Resid. d.f.* | *Resid. Dev.* | *F* | *p* |
|---|---|---|---|---|---|---|
| Null Model | - | - | 97 | 83.94 | - | - |
| Mycorrhizal Inoculation | 1 | 9.16 | 96 | 74.78 | 9.16 | **0.003** |
| Field Epidemic Population | 2 | 3.08 | 94 | 71.70 | 1.54 | 0.21 |
| Seed Origin Population | 5 | 6.94 | 89 | 64.76 | 1.39 | 0.22 |
| Host Size | 1 | 1.04 | 88 | 63.72 | 1.04 | 0.31 |

**Table S12** Factors influencing the proportion of infected leaves in hosts in thirty maternal genotypes at the end of the field epidemic experiment. We examined the effects of inoculation with arbuscular mycorrhizal fungi, maternal genotype, field epidemic population, host size (Table S12B) and their interactions on the proportion of infected leaves in *Pl. lanceolata* individuals at the end of the field epidemic experiment using a generalized logistic regression model ($n$ = 210 individuals). Non-significant interaction terms were removed from the final model. Table S12B shows the results of the model with host initial size included. Analysis of deviance results are shown in the table below, including degrees of freedom (d.f.), deviance, residual d.f., residual deviance, $F$-score, and $p$-values. Effects for which $p < 0.05$ are highlighted in bold.

**Table S12A: Without Host Size**

| Analysis of Deviance | *d.f.* | *Deviance* | *Resid. d.f.* | *Resid. Dev.* | *F* | *p* |
|---|---|---|---|---|---|---|
| Null Model | - | - | 97 | 83.94 | - | - |
| Mycorrhizal Inoculation | 1 | 9.16 | 96 | 74.78 | 9.16 | **0.002** |
| Field Epidemic Population | 2 | 3.08 | 94 | 71.70 | 1.54 | 0.21 |
| Maternal Genotype | 28 | 36.77 | 66 | 34.94 | 1.31 | 0.12 |

**Table S12B: With Host Size**

| Analysis of Deviance | *d.f.* | *Deviance* | *Resid. d.f.* | *Resid. Dev.* | *F* | *p* |
|---|---|---|---|---|---|---|
| Null Model | - | - | 97 | 83.94 | - | - |
| Mycorrhizal Inoculation | 1 | 9.16 | 96 | 74.78 | 9.16 | **0.002** |
| Field Epidemic Population | 2 | 3.08 | 94 | 71.70 | 1.54 | 0.21 |
| Maternal Genotype | 28 | 36.77 | 66 | 34.94 | 1.31 | 0.12 |
| Host Size | 1 | 0.78 | 65 | 34.16 | 0.78 | 0.38 |

**Table S13** Factors influencing the number of infected leaves in hosts in six seed origin populations during the epidemic peak. We examined the effects of inoculation with arbuscular mycorrhizal fungi, seed origin population, field epidemic population, host size (Table S13B), and their interactions on the number of infected leaves in *Pl. lanceolata* individuals during the peak of the field epidemic experiment using a generalized logistic regression model ($n$ = 210 individuals). Non-significant interaction terms were removed from the final model. Table S13B shows the results of the model with host initial size included. Analysis of deviance results are shown in the table below, including degrees of freedom (d.f.), deviance, residual d.f., residual deviance, $F$-score, and $p$-values. Effects for which $p < 0.05$ are highlighted in bold.

#### Table S13A: Without Host Size

| Analysis of Deviance | *d.f.* | *Deviance* | *Resid. d.f.* | *Resid. Dev.* | *F* | *p* |
|---|---|---|---|---|---|---|
| Null Model | - | - | 209 | 3981.3 | - | - |
| Mycorrhizal Inoculation | 1 | 0.98 | 208 | 3980.3 | 0.05 | 0.82 |
| Seed Origin Population | 5 | 320.11 | 203 | 3660.2 | 3.43 | **0.005** |
| Field Epidemic Population | 2 | 82.08 | 201 | 3578.1 | 2.20 | 0.11 |

#### Table S13B: With Host Size

| Analysis of Deviance | *d.f.* | *Deviance* | *Resid. d.f.* | *Resid. Dev.* | *F* | *p* |
|---|---|---|---|---|---|---|
| Null Model | - | - | 209 | 3981.3 | - | - |
| Mycorrhizal Inoculation | 1 | 0.98 | 208 | 3980.3 | 0.05 | 0.82 |
| Seed Origin Population | 5 | 320.11 | 203 | 3660.2 | 3.43 | **0.005** |
| Field Epidemic Population | 2 | 82.08 | 201 | 3578.1 | 2.20 | 0.11 |
| Host Size | 1 | 106.92 | 200 | 3471.2 | 5.81 | **0.01** |

**Table S14** Factors influencing the number of infected leaves in hosts in thirty maternal genotypes at the peak of the field epidemic experiment. We examined the effects of inoculation with arbuscular mycorrhizal fungi, maternal genotype, field epidemic population, host size (Table S14B), and their interactions on the proportion of infected leaves in *Pl. lanceolata* individuals at the peak of the field epidemic experiment using a generalized logistic regression model ($n$ = 210 individuals). Host size was included as the number of leaves on each individual at the peak of the epidemic in Table S14B. Non-significant interaction terms were removed from the final model. Analysis of deviance results are shown in the table below, including degrees of freedom (d.f.), deviance, residual d.f., residual deviance, $F$-score, and $p$-values. Effects for which $p < 0.05$ are highlighted in bold.

**Table S14A: Without Host Size**

| Analysis of Deviance | *d.f.* | *Deviance* | *Resid. d.f.* | *Resid. Dev.* | *F* | *p* |
|---|---|---|---|---|---|---|
| Null Model | - | - | 209 | 3981.3 | - | - |
| Mycorrhizal Inoculation | 1 | 0.98 | 208 | 3980.3 | 0.08 | 0.77 |
| Maternal Genotype | 29 | 959.35 | 179 | 3021.0 | 2.80 | **1.01 x $e^{-04}$** |
| Field Epidemic Population | 2 | 115.85 | 177 | 2905.1 | 4.90 | **0.01** |
| Myc. Inoc. × Mat. Gen. | 29 | 545.57 | 148 | 2359.5 | 1.59 | **0.05** |
| Mat. Gen. × Field Pop. | 55 | 1217.22 | 93 | 1142.3 | 1.87 | **0.004** |

**Table S14B: With Host Size**

| Analysis of Deviance | *d.f.* | *Deviance* | *Resid. d.f.* | *Resid. Dev.* | *F* | *p* |
|---|---|---|---|---|---|---|
| Null Model | - | - | 209 | 3981.3 | - | - |
| Mycorrhizal Inoculation | 1 | 0.98 | 208 | 3980.3 | 0.09 | 0.77 |
| Maternal Genotype | 29 | 959.35 | 179 | 3021.0 | 1.96 | **0.004** |
| Field Epidemic Population | 2 | 115.85 | 177 | 2905.1 | 3.43 | **0.03** |
| Host Size | 1 | 106.63 | 176 | 2798.5 | 6.31 | **0.01** |

**Table S15** Factors influencing the number of infected leaves in hosts in six seed origin populations at the end of the field epidemic experiment. We examined the effects of inoculation with arbuscular mycorrhizal fungi, seed origin population, field epidemic population, host size (Table S15B), and their interactions on the number of infected leaves in *Pl. lanceolata* individuals at the end of the field epidemic experiment using a generalized logistic regression model ($n$ = 98 individuals). Host size was included as the number of leaves on each individual at the end of the epidemic. Table S15B shows the results of the model with host initial size included. Non-significant interaction terms were removed from the final model. Analysis of deviance results are shown in the table below, including degrees of freedom (d.f.), deviance, residual d.f., residual deviance, $F$-score, and $p$-values. Effects for which $p < 0.05$ are highlighted in bold.

#### Table S15A: Without Host Size

| Analysis of Deviance | *d.f.* | *Deviance* | *Resid. d.f.* | *Resid. Dev.* | *F* | *p* |
|---|---|---|---|---|---|---|
| Null Model | - | - | 97 | 1778.1 | - | - |
| Mycorrhizal Inoculation | 1 | 51.88 | 96 | 1726.2 | 2.57 | 0.11 |
| Seed Origin Population | 5 | 205.14 | 91 | 1521.0 | 2.03 | 0.08 |
| Field Epidemic Population | 2 | 164.72 | 89 | 1356.3 | 4.09 | **0.02** |

#### Table S15B: With Host Size

| Analysis of Deviance | *d.f.* | *Deviance* | *Resid. d.f.* | *Resid. Dev.* | *F* | *p* |
|---|---|---|---|---|---|---|
| Null Model | - | - | 97 | 1778.1 | - | - |
| Mycorrhizal Inoculation | 1 | 51.88 | 96 | 1726.2 | 2.99 | 0.09 |
| Seed Origin Population | 5 | 205.14 | 91 | 1521.0 | 2.37 | **0.05** |
| Field Epidemic Population | 2 | 164.72 | 89 | 1356.3 | 4.75 | **0.01** |
| Host Size | 1 | 185.71 | 88 | 1170.6 | 10.71 | **0.002** |

**Table S16** Factors influencing the number of infected leaves in hosts in thirty maternal genotypes at the end of the field epidemic experiment. We examined the effects of inoculation with arbuscular mycorrhizal fungi, maternal genotype, field epidemic population, host size (Table S16B), and their interactions on the number of infected leaves in *Pl. lanceolata* individuals at the end of the field epidemic experiment using a generalized logistic regression model (*n* = 98 individuals). Host size was included as the number of leaves on each individual at the end of the epidemic. Table S16B shows the results of the model with host initial size included. Non-significant interaction terms were removed from the final model. Analysis of deviance results are shown in the table below, including degrees of freedom (d.f.), deviance, residual d.f., residual deviance, *F*-score, and *p*-values. Effects for which $p < 0.05$ are highlighted in bold.

#### Table S16A: Without Host Size

| **Analysis of Deviance** | ***d.f.*** | ***Deviance*** | ***Resid. d.f.*** | ***Resid. Dev.*** | ***F*** | ***p*** |
|---|---|---|---|---|---|---|
| Null Model | - | - | 97 | 1778.06 | - | - |
| Mycorrhizal Inoculation | 1 | 51.88 | 96 | 1726.17 | 4.51 | **0.04** |
| Maternal Genotype | 28 | 821.89 | 68 | 904.29 | 2.55 | **0.001** |
| Field Epidemic Population | 2 | 150.45 | 66 | 753.83 | 6.54 | **0.003** |

#### Table S16B: With Host Size

| **Analysis of Deviance** | ***d.f.*** | ***Deviance*** | ***Resid. d.f.*** | ***Resid. Dev.*** | ***F*** | ***p*** |
|---|---|---|---|---|---|---|
| Null Model | - | - | 97 | 1778.06 | - | - |
| Mycorrhizal Inoculation | 1 | 51.88 | 96 | 1726.17 | 4.79 | **0.03** |
| Maternal Genotype | 28 | 821.89 | 68 | 904.29 | 2.71 | **4.92 x $e^{-04}$** |
| Field Epidemic Population | 2 | 150.45 | 66 | 753.83 | 6.94 | **0.002** |
| Host Size | 1 | 74.17 | 65 | 679.67 | 6.85 | **0.01** |

**Table S17** Relationship between host growth and defensive effects due to mycorrhizal inoculation in the maternal genotypes. We examined the relationship between the growth and defensive effects following inoculation with arbuscular mycorrhizal fungi in individuals of 30 maternal genotypes using a linear regression model. Defensive effects were included as the response variable and are examined at the peak of the epidemic using the change in the number of infected leaves (in infected individuals) when the genotype was associated with mycorrhizal fungi. Specifically, defensive effects due to mycorrhizal inoculation are quantified as the effect size resulting from a model contrasting infection in mycorrhizal vs. non-mycorrhizal plants within each genotype. Growth effects were quantified similarly, using the effect size coefficients resulting from a model contrasting host growth (changes in the total number of leaves) in mycorrhizal vs. non-mycorrhizal plants within each genotype at the beginning of the experiment. Analysis of variance (ANOVA) results are shown in the table below, including degrees of freedom (d.f.), sum of squares (SS), mean square (MS), *F*-statistic, and *p*-value for the explanatory factor.

| ANOVA | *d.f.* | *SS* | *MS* | *F* | *p* |
|---|---|---|---|---|---|
| Myc. Effect on Host Growth | 1 | 4.40 | 4.40 | 2.76 | 0.11 |
| Residuals | 28 | 44.52 | 1.59 | - | - |

**Table S18** Relationship between disease susceptibility and defensive effects due to mycorrhizal inoculation in the maternal genotypes. We examined the relationship between baseline disease susceptibility and the defensive effects of arbuscular mycorrhizal inoculation in individuals of 30 maternal genotypes using a linear regression model. Defensive effects are included as the response variable and are examined at the peak of the epidemic using the change in the number of infected leaves (in infected individuals) when the genotype was associated with mycorrhizal fungi. Specifically, defensive effects due to mycorrhizal inoculation are quantified as the effect size resulting from a model contrasting infection in mycorrhizal vs. non-mycorrhizal plants within each genotype. Disease susceptibility is quantified for each maternal genotype as the estimated coefficient representing the number of infected leaves in infected, non-mycorrhizal individuals. Analysis of variance (ANOVA) results are shown in the table below, including degrees of freedom (d.f.), sum of squares (SS), mean square (MS), *F*-statistic, and p-value for the explanatory factor. Effects for which $p < 0.05$ are highlighted in bold.

| ANOVA | *d.f.* | *SS* | *MS* | *F* | *p* |
|---|---|---|---|---|---|
| Disease Susceptibility | 1 | 22.03 | 22.03 | 22.94 | **4.93 x e$^{-05}$** |
| Residuals | 28 | 26.89 | 0.96 | - | - |
